# The PP1 phosphatase complex coordinates pre-mRNA 3’end processing and termination of RNA polymerase II transcription

**DOI:** 10.64898/2025.12.17.694969

**Authors:** Alex Au, Abigail Southers, Krzysztof Kuś, Niek Schoemaker, Esra Balikci-Akil, Charles Hewes, Marjorie Fournier, Ebru Aydin, Sarah Sayed Hassanein, Cornelia Kilchert, Jonathan M. Grimes, Lidia Vasiljeva

## Abstract

Phosphorylation plays a central role in coordinating transcription with pre-mRNA maturation, yet the mechanistic understanding of how transcription is regulated by phosphorylation remains limited. The PP1 phosphatase removes CDK9-dependent phosphorylation of Pol II and Spt5, triggering transcription termination. Here, we demonstrate that PNUTS enhances PP1 activity, whereas WDR82 mediates an interaction between PP1 and the pre-mRNA 3′ end processing machinery subunit Symplekin. We present the first structure of a PNUTS–WDR82–Symplekin–Ssu72 complex, revealing that PNUTS and Symplekin bind WDR82 through distinct interfaces using conserved short linear motifs. Mutations that inactivate PP1’s catalytic activity or disrupt its interaction with either PNUTS or the 3′ end processing machinery led to impaired transcription elongation, inefficient cleavage of nascent transcripts, altered poly(A) site selection, and widespread transcription termination defects. Our findings demonstrate that the PP1 complex is essential for coupling transcription to pre-mRNA 3′ end maturation, ensuring production of functional mRNA.

## INTRODUCTION

Transcription of protein-coding genes by RNA Polymerase II (Pol II) is a tightly regulated process that is divided into three stages: initiation, elongation, and termination. Eukaryotic mRNA precursors (pre-mRNAs) must undergo extensive processing, including 5’-capping, splicing, 3’end cleavage and polyadenylation, which are essential to produce a functional mRNA. Coupling of mRNA processing to transcription is important to control the accuracy, efficiency and order of mRNA processing reactions. Dynamic phosphorylation of the intrinsically disordered carboxy-terminal domain (CTD) of the Pol II Rpb1 subunit, and the conserved transcription elongation factor Spt5 plays important role in co-transcriptional RNA processing^1–4^. The CTD is composed of the Y_1_S_2_P_3_T_4_S_5_P_6_S_7_ heptad repeats, within which Tyrosine 1 (Tyr1), Serine 2 (Ser2), Threonine 4 (Thr4), Serine 5 (Ser5) and Serine 7 (Ser7) are reversibly phosphorylated during the transcription cycle^5,6^. Kinases and phosphatases coordinate the addition and removal of phosphate groups, generating phosphorylation patterns characteristic for each transcriptional stage, which enables temporal recruitment of RNA processing factors.

Spt5 forms the DRB (5,6-Dichlorobenzimidazole riboside) Sensitivity-Inducing Factor (DSIF) complex with Spt4^7–9^. DSIF is recruited to Pol II during early elongation and remains associated with Pol II throughout the transcription cycle. Unphosphorylated Spt5 stabilises Pol II promoter-proximal pausing. Release of Pol II from promoter proximal pausing into elongation is driven by the Cdk9 kinase of the P-TEFb complex. Cdk9 phosphorylates Spt5, Ser2 of the Pol II CTD, and the Negative Elongation Factor (NELF). Phosphorylation of Spt5 is associated with the productive elongation, whereas loss of Spt5 phosphorylation impairs elongation and leads to termination of transcription^10–13^. In fission yeast, *Schizosaccharomyces pombe* (*S. pombe*), Spt5 contains 18 tandem repeats in its C-terminal region (CTR) that are phosphorylated at Thr1 within the consensus motif T₁P₂A₃W₄N₅T₆G₇R₈/K. In mammalian cells, CTR is phosphorylated on Thr4 within the consensus repeat G_1_S_2_Q/R_3_T_4_P_5_X_6_Y_7_, as well as within a flexible linker between KOW4 and 5 domains^12,13^. Although Spt5 phosphorylation is essential for Pol II processivity, its precise mechanistic contributions remain unclear. Similar to the Pol II CTD, the Spt5 CTR likely functions as a phosphorylation-dependent scaffold that coordinates the recruitment of elongation factors such as the PAF1 complex subunit Rtf1, which promotes productive elongation^14–17^.

In eukaryotes, Pol II dislodgement and transcription termination are mechanistically linked to the endonucleolytic cleavage of nascent RNA^18–22^. At the 3’ end of the gene, nascent RNA is cleaved by the multi-subunit *<u>C</u>*leavage and *<u>P</u>*oly*<u>A</u>*denylation machinery (CPA), which also adds polyadenylate tail to the 3’end of mRNA. CPA is composed of the Cleavage and Polyadenylation Factor (CPF) and Cleavage Factor IA (CFIA), as well as the Cleavage Factor IB (CFIB) in yeast and Cleavage and Polyadenylation Specificity Factor (CPSF), Cleavage Factors (mCF) I and II and cleavage stimulation factor complex (CstF) in mammalian cells^18^. The CPF (CPSF in mammals) recognises the AAUAAA element of the polyadenylation signal (PAS). The cleavage factors recognize auxiliary sequence elements around the polyadenylation signal (PAS), which help define the site of the 3′ end processing in pre-mRNA^23–28^. The endonuclease subunit of the CPA (Ysh1 in yeast and CPSF73 in mammals) cleaves the nascent RNA, generating a 5’-monophosphorylated end that is targeted by the transcription termination factor - 5’ to 3’ exoribonuclease Xrn2^29–31^. Xrn2-mediated degradation of nascent RNA is important for the dislodgement of Pol II from DNA^29,32^.

Accurate selection of PAS by the CPA determines the position of the cleavage, lengths and composition of the 3’UTR that can affect mRNA localisation, translation efficiency and stability ^33–35^. Recruitment of CPA to PAS within the gene body leads to premature transcription termination antagonising production of mRNA and attenuating the expression of selected protein-coding genes^20,22,36–40^. Yet, the mechanisms underlying PAS selection by the CPA are not well understood.

In addition to CPA, there is another large endonucleolytic complex, called Integrator, which is only present in metazoans. Integrator is recruited by the DSIF to the NELF-bound paused elongation complex to enable premature transcription termination^20–22,41,42^.

While phosphorylated Spt5 acts as a positive elongation factor, dephosphorylation of Spt5 by the conserved PP1 is critical for transcription termination in both yeast and metazoans^12,43–46^. Removal of phosphorylation is associated with deceleration of Pol II downstream of the PAS, which was proposed to provide kinetic advantage to Xrn2 as it degrades nascent RNA emerging from polymerase to mediate Pol II dislodgement from DNA^46^. However, recent work has demonstrated that Xrn2 forms a stable complex with Pol II and requires Spt5 for enhancing its nucleolytic activity, which is essential for Pol II dislodgement^32,47,48^. These findings suggest that Spt5 dephosphorylation could contribute to the other aspects of this process such as PAS choice and/or cleavage efficiency of the nascent transcript by the CPA.

PP1 phosphatase also dephosphorylates the Pol II CTD. In yeast, PP1 dephosphorylates the CTD consensus phosphorylated on Thr4 and Tyr1, whereas in mammalian cells, it primarily targets phosphorylated Ser5 ^43,44,49–52^. However, the mechanisms by which non-specific phosphatases such as PP1 are recruited and directed to act in a context-dependent manner during transcription, including termination, remain poorly understood. Physiologically relevant PP1 holoenzymes form through the association of catalytic and regulatory subunits^53–58^. Different regulatory subunits assemble in PP1 complexes with distinct substrate specificities that function in various cellular pathways^57,59–61^. PP1 interacts with its regulatory subunits through short linear motifs (SLiMs) that dock to specific surface grooves on the catalytic subunit. The most prevalent is the hydrophobic RVxF motif, which anchors the regulatory subunit to PP1^53,54,62,63^. The PP1-Nuclear-Targeting Subunit (PNUTS) and the WD40-repeat protein WDR82 interact with and have been proposed to mediate the function of nuclear PP1 in transcription in mammals^46,49,64^. In addition to Spt5 and Pol II, PNUTS has been linked to the regulation of PP1-mediated dephosphorylation of other proteins such as MYC, p53 and the retinoblastoma protein^65–68^. Depletion of PNUTS and WDR82 in mammalian cells leads to failed transcription termination, resulting in read-through transcription at the end of transcription units^49^. More recently, depletion of PNUTS and WDR82 have been shown to affect promoter-proximal pause release and to prevent premature transcription termination at promoter-proximal regions and gene enhancers^51,52,69,70^.

Notably, PP1 and its putative regulatory subunits have been co-purified with the 3’end processing machinery in both budding yeast *Saccharomyces cerevisiae* (*S. cerevisiae*) and *S. pombe*^43,71–73^. In budding yeast, PP1 (Glc7) forms part of the APT complex (Associated with Pta1) together with Ref2 (*S. cerevisiae* homologue of PNUTS), SWD2 (*S. cerevisiae* homologue of WDR82), the CPA scaffolding subunit Pta1/Symplekin and the Ser5 phosphatase Ssu72^74–78^. The APT complex has been functionally linked to the cleavage-independent 3′ end formation and transcription termination of non-coding RNAs and suppression of cryptic transcription together with the trimeric RNA binding Nrd1-Nab3-Sen1 termination complex (NNS)^79–83^. In metazoans, a functionally analogous pathway relies on the Restrictor complex, composed of arginine-rich zinc-finger protein ZC3H4 (mammals)/SU(S) (Suppressor of Sable) (*Drosophila*) and WDR82^70,84,85^, which cooperates with PP1–PNUTS to suppress cryptic transcription and promote premature termination^70,84–88^. Despite this progress revealing conserved role of PP1 in transcription, how PP1 holoenzymes are organised and activated on chromatin to regulate this process remains unclear.

Our study discovers a key role for the conserved PP1–PNUTS–WDR82 complex in coupling Pol II transcription to pre-mRNA 3′ end maturation. Here, using *S. pombe* as a model, we reconstitute a trimeric PP1–PNUTS–WDR82 complex and show that PNUTS stimulates PP1 activity toward Spt5, whereas WDR82 connects the phosphatase to the 3′ end processing machinery through CPA subunit Symplekin. We delineate the molecular principles that regulate PP1 activity, assembly of the functional complex and its co-transcriptional tethering. Our nascent transcriptome analyses revealed that loss of PP1 activity, disruption of its interaction with PNUTS, or perturbation of PP1 holoenzyme tethering to Symplekin leads to altered PAS choice, defective 3′ end cleavage, transcriptional readthrough and impaired elongation. We propose that multivalent interactions mediated by WDR82 constitute a combinatorial code that enables crosstalk among multiple complexes to regulate transcription.

## RESULTS

### PNUTS/Ppn1 and WDR82/Swd2.2 form a stable complex with PP1/Dis2

PNUTS and WDR82 are conserved across eukaryotes, with the fission yeast homologs Ppn1 and Swd22 showing slightly higher identity to their human counterparts (global alignment 12.3% and 33.2%, respectively) than the corresponding budding yeast proteins (10.9% and 31.9%) (Figure 1A). The genetic tractability of yeast makes it an ideal organism to explore the mechanistic role of PP1 and its regulatory subunits in transcription. For simplicity, we refer to the *S. pombe* orthologues by the metazoan names in this study. Like mammalian PNUTS, the *S. pombe* orthologue is exclusively nuclear, which suggests its role in targeting PP1 to nuclear substrates is conserved (Figure S1A). Secondary structure predictions suggest that PNUTS is largely unstructured, except for its N-terminal region, which adopts a HEAT domain followed by a TFIIS N-terminal–domain–like (TND-like) fold. The remaining disordered regions contain several linear motifs, including a predicted PP1-interacting RvXF motif located at residues 506–510 of *S. pombe* PNUTS (Figure 1A).

**Figure 1.**
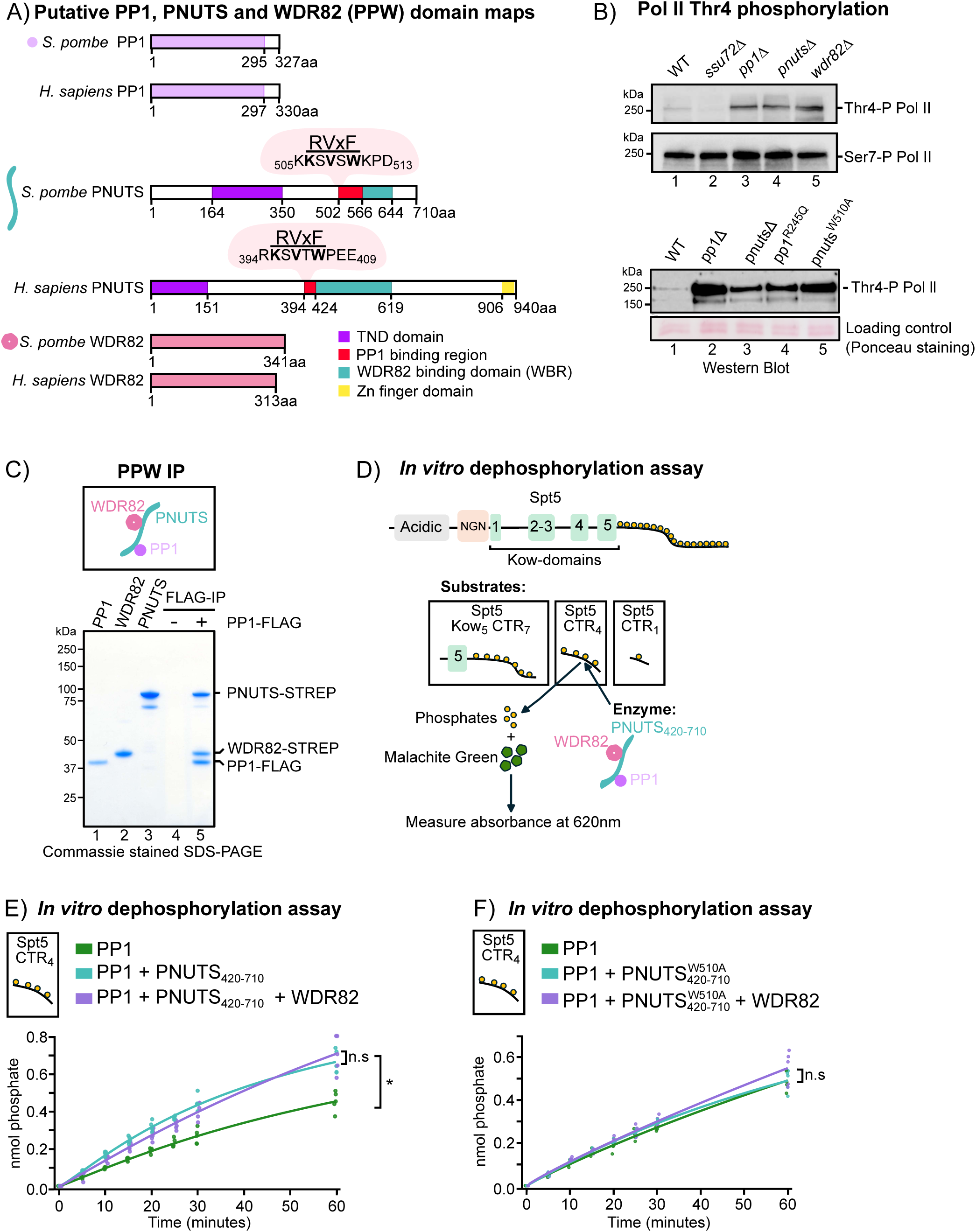
A) Domain maps of PP1, PNUTS, and WDR82 from *S. pombe* and *H. sapiens.* TND-TFIIS-N-terminal domain. B) Western blots to assess levels of Thr4-P Pol II were performed in PP1 complex subunit perturbations including *pp1Δ, pnutsΔ, wdr82Δ, pp1*^R245Q^, *pnuts*^W510A^. WT and *ssu72Δ* strains were used as controls. Ponceau staining of the membrane and CTD Ser7-P levels were used as a loading control. C) Reconstitution of PPW trimeric complex. Purified PP1–FLAG, WDR82, and PNUTS were loaded as inputs. WDR82 and PNUTS co-immunoprecipitated with PP1–FLAG on M2 beads. Beads were incubated with WDR82-PNUTS as negative control. D) Schematic depiction of the phosphatase assay and overview of the Spt5 truncation construct and phosphorylated peptides used as substrates. E) Phosphatase assays were performed using PP1 alone, PP1 with PNUTS, or PP1 with PNUTS and WDR82, using a 4× phosphorylated peptide substrate (Spt5-CTR_4_). PNUTS enhances PP1-mediated dephosphorylation of Spt5-CTR_4_, whereas WDR82 does not display a similar stimulatory effect. F) PNUTS_420-710_^W510A^ cannot enhance PP1-mediated dephosphorylation of Spt5-CTR_4._

To explore whether PNUTS and WDR82 are required for PP1 activity, we assessed the phosphorylation status of known PP1 substrates in whole cell extract^43^. Analysis of the Pol II Thr4-P levels, which is a key modification universally associated with transcription termination^43,89–91^ revealed that loss of either PNUTS or WDR82 leads to dramatic accumulation of hyper-phosphorylated Thr4 form of Pol II (Figure 1B, top panel lanes 4-5 and bottom panel lane 3), which was similar to what is observed in cells lacking active PP1 (PP1^R245Q^ mutant) (Figure 1B, lane 4, bottom panel). In contrast, no increase in Thr4-P was observed upon loss of Ssu72, which is Pol II Ser5-P specific phosphatase^76,77^ (Figure 1B, lane 2, upper panel). These results suggest that PP1 relies on PNUTS and WDR82 for dephosphorylation of its substrates during Pol II transcription. Consistent with this hypothesis, mutating the tryptophan residue to an alanine in the _506_KSVSW_510_ portion of the PNUTS RvXF motif predicted to abolish PNUTS-PP1 interaction (PNUTS^W510A^) leads to increased accumulation of hyper-phosphorylated Pol II Thr4-P (Figure 1B, bottom panel lane 5), further supporting the importance of the PNUTS-PP1 interaction in dephosphorylating PP1 substrates during Pol II transcription.

To understand the roles of PNUTS and WDR82 in PP1-mediated dephosphorylation during Pol II transcription, we undertook a reductionist *in vitro* approach using purified proteins. First, we tested whether PP1, PNUTS and WDR82 can form a trimeric complex by incubating individually expressed and purified proteins (Figure 1C, lanes 2 and 3). SDS-PAGE analyses revealed the presence of PNUTS and WDR82 in PP1-FLAG elution, whereas no proteins were recovered from control IP lacking PP1-FLAG (Figure 1C, lanes 4 and 5). We conclude that the formation of a trimeric PP1-PNUTS-WDR82 (PPW) complex is evolutionarily conserved^51,52,64^.

### PNUTS stimulates PP1 activity against Spt5-CTR

To gain insight into how PNUTS and WDR82 contribute to PP1 phosphatase activity, we established an *in vitro* dephosphorylation assay (Figure 1D). PNUTS directly binds WDR82 and PP1 via distinct and adjacent binding regions (Figure 1A)^71^. Since the N-terminal part of PNUTS was shown to be dispensable for interactions with PP1 and WDR82^51,64^, we expressed and purified recombinant PNUTS 420-720 harbouring the PP1-interacting RvXF motif and a region predicted to interact with WDR82^64,71^ (Figure 1A), WDR82 and PP1 (Figure S1B). To determine the contribution of the PPW subunits to PP1 activity, we tested its activity *in vitro* against various Spt5 substrates. Firstly, a phosphorylated Spt5 KOW_5_ construct containing seven CTR consensus repeats (Figure S1C), then a synthetic phospho-Spt5 peptides comprising four and one repeats of the consensus CTR (Figure S1D). Phosphatase activity assay was conducted either in the presence or absence of PNUTS and WDR82 and then quantified by measuring phosphate release over time (Figure 1D). Interestingly, the addition of PNUTS stimulated PP1 activity 1.5-fold for all the substrates (Figures 1E and S1E and F). Notably, PP1 activity and PNUTS-dependent stimulation were comparable for all the substrates tested suggesting that one repeat peptide is sufficient for the enzyme to engage with the substrate (Figure 1E, F, S1E and F). In contrast, the PNUTS^W510A^ mutant failed to stimulate PP1 activity (Figures 1F), indicating that the RVxF motif-mediated interaction between PNUTS and PP1 is critical for PNUTS-dependent regulation of PP1 activity. Surprisingly, the addition of WDR82 does not stimulate PP1 activity with or without functional PNUTS present (Figure 1E, F) suggesting that WDR82 is important for other aspects of PP1-mediated dephosphorylation. Together, these results indicate that PNUTS enhances PP1 phosphatase activity towards Spt5 CTR, whereas WDR82 does not contribute to PP1 activity stimulation.

### Loss of PP1 activity or regulatory subunits leads to altered pre-mRNA 3’end processing, and defective transcription termination

To investigate the roles of PP1, PNUTS and WDR82 in transcription, we generated mutant strains to dissect their regulatory and catalytic contributions *in vivo.* We created a strain where PNUTS cannot bind and activate PP1 (*pnuts*^W510A^, in which the endogenous protein is complemented by a mutant allele in the *leu* locus) (Figure 1A and F), alongside a catalytically inactive *pp1*^R245Q^ mutant that is unable to dephosphorylate PP1 substrates^92^ (Figure 1B).

To directly monitor nascent transcription, we performed spike-in normalised transient transcriptome sequencing (TT-seq) in *pp1*Δ, and *pp1*^R245Q^*, wdr82*Δ, *pnuts*Δ and *pnuts*^W510A^ strains. TT-seq revealed a genome-wide elongation defect in all PPW mutants, leading to reduced expression of protein-coding genes (Figures 2A and S2A). While non-coding RNAs were less affected across mutants, ribosomal RNA genes showed modest downregulation specifically in *pp1*Δ and *pp1*^R245Q^ strains (Figure S2B). Strikingly, all mutants exhibited elevated reads downstream of annotated polyadenylation sites (PAS), as demonstrated at individual gene loci (Figure 2B) and in genome-wide analyses normalised to gene expression (Figure 2C), indicating defective 3’-end processing and transcription termination. Additionally, the PPW mutants exhibited alternative polyadenylation (APA), with many genes shifting to more distal poly(A) sites (Figure 2D). To analyse alternative polyadenylation, we examined genes with 3’UTRs ≥50nt and identified those with >1.5-fold increase in distal PAS usage relative to WT in at least two mutants, revealing 514 genes with significantly longer 3’UTRs (3419 unchanged genes) (Figure S2C). The upregulated 3’UTRs observed in PPW mutants exhibited a distinctive bimodal GC content distribution with a second peak at higher GC content (Figure S2D). Furthermore, hexamer analysis revealed enrichment of CG-rich motifs in genes with a shift in 3’UTR usage (Figure S2E), suggesting that the dependency of 3′-end processing on PPW may be modulated by 3’UTR sequence context. We speculate that sequence could either contribute to affinity of CPA binding to nascent RNA or could potentially affect speed of Pol II or pausing frequency. Together, this data demonstrated that the PP1–PNUTS–WDR82 phosphatase complex is essential for efficient transcription elongation, accurate 3′-end formation, and proper termination. Loss of PP1 catalytic activity or disruption of its regulatory subunits induces genome-wide elongation defects and pervasive readthrough transcription. The PP1 complex acts as a key integrator coupling transcription elongation with RNA 3′-end processing and termination. We propose that the PPW complex regulates CPA-mediated PAS selection.

**Figure 2.**
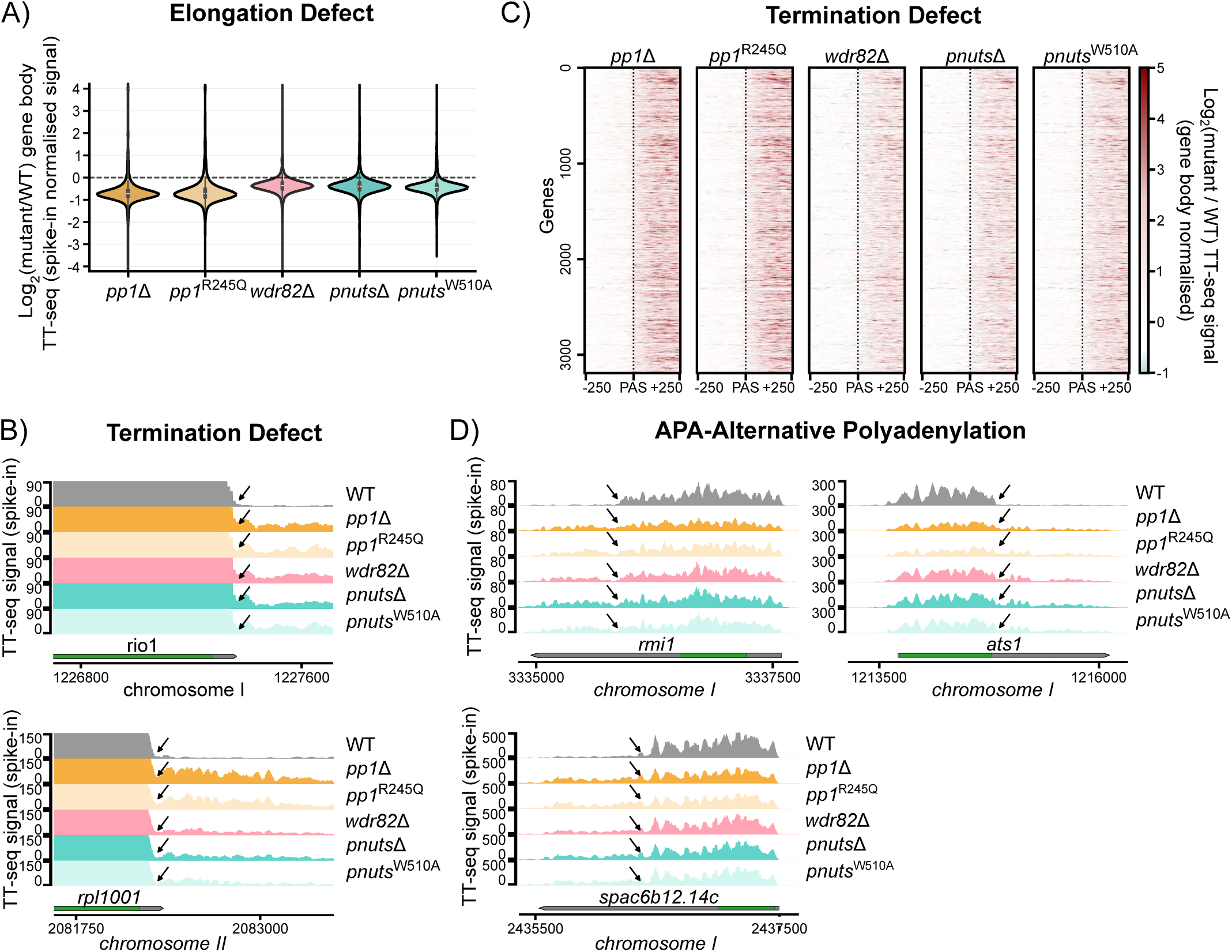
A) Relative changes in nascent transcription upon loss of PP1, its catalytic activity or regulatory subunits (WDR82, PNUTS or complementation strain with PNUTS^W510A^ which losses interaction with PP1). Expression levels across protein coding genes were measured by the TT-seq spike-in normalised and referenced to the WT strain. B) Transcriptional readthrough which is a potential hallmark of transcription termination defects for representative genes in mutants described in (A). C) Heatmaps representing termination defects (readthrough) scaled to expression in gene body are global and affect vast majority of protein coding genes. D) Mutants described in (A) show changes in 3’UTR usage. Arrows highlight proximal PAS used predominantly in WT cells.

### WDR82 links PNUTS-PP1 to the CPA machinery

We next asked how the PPW complex participates in mRNA 3’end processing. To gain insight into PPW function, we purified *S. pombe* FTP-tagged PNUTS using tandem affinity chromatography and identified proteins co-purified with PNUTS by mass spectrometry (Figure 3A). As is the case for *S. cerevisiae* and consistent with previously published work, the PPW complex in *S. pombe* co-purifies with the CPA machinery (Figure 3A)^73^. In *S. cerevisiae*, the PPW complex interacts with additional budding yeast specific subunits Syc1 and Pti1 and forms a complex with Pta1 (Symplekin) and Ssu72, which was termed the Phosphatase Module within CPA^93,94^. Although it is not fully clear how PPW integrates within *S. cerevisiae* CPF, noncovalent nanoelectrospray ionization mass spectrometry (nanoESI-MS) combined with computational analysis suggested that the PPW complex interacts with Syc1 and Pti1, which could tether PPW to CPF via Symplekin^94^. We therefore hypothesised that PPW forms a complex with the 3’end processing machinery. Indeed, a stable 12-subunit complex between the *S. pombe* CPF and the PP1 holoenzyme can be reconstituted *in vitro* using a baculovirus expression system (Figure 3B). This indicates that the PP1 holoenzyme indeed is a part of the CPA machinery, which may explain why defective 3′-end processing is observed by TT-seq upon loss of PPW subunits (Figure 2D, S2C-E).

**Figure 3.**
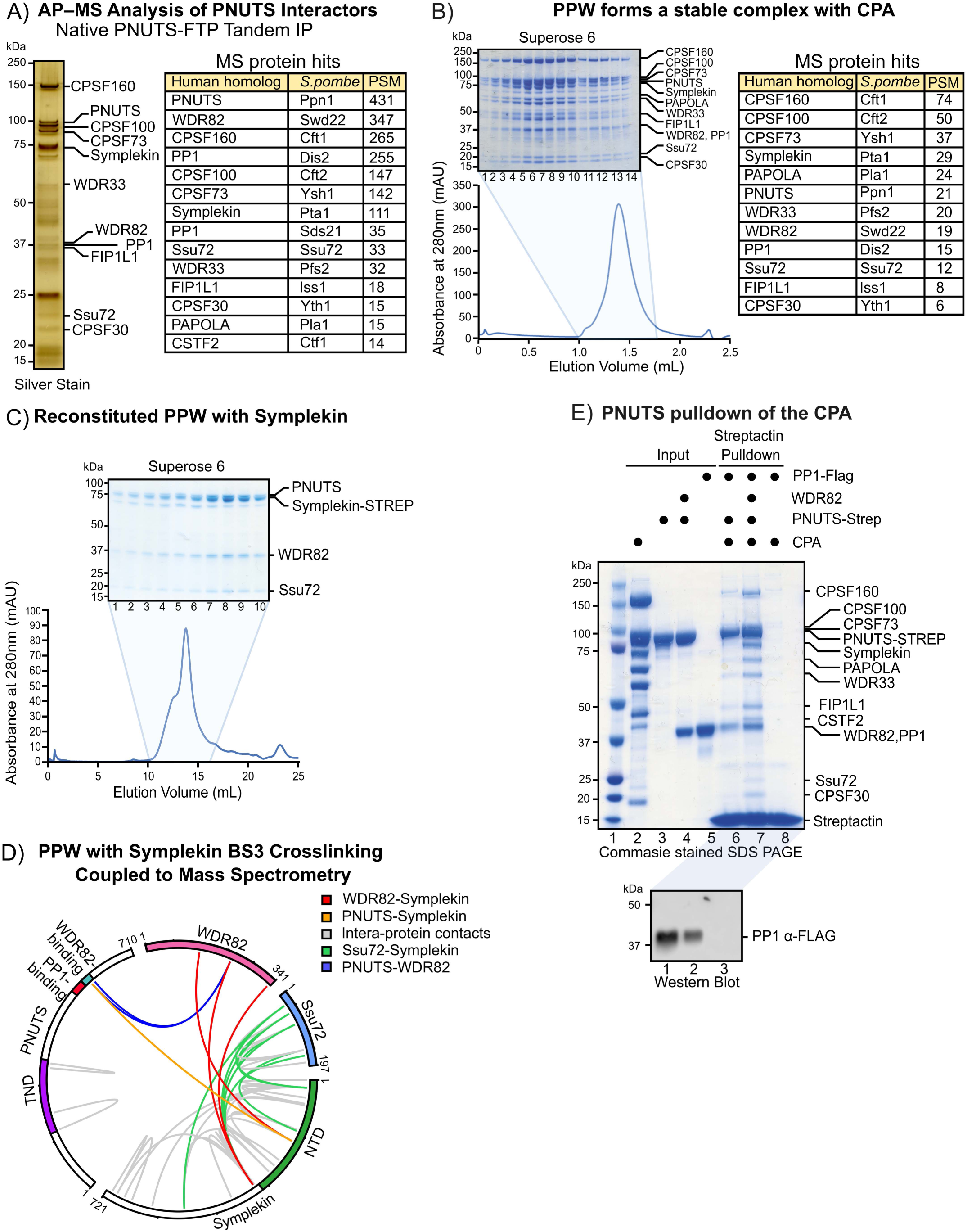
A) FTP-tagged PNUTS was purified from *S. pombe*, and proteomic analysis of co-purifying factors identified PP1, WDR82, and multiple components of the cleavage and polyadenylation (CPA) machinery. B) A stable 12-subunit complex comprising the *S. pombe* CPA machinery and the PP1 holoenzyme was reconstituted *in vitro*. Table shows the results of the mass spectrometry analyses. C) Biochemical reconstitution demonstrates that PNUTS, WDR82, Symplekin, and Ssu72 assemble as a stable tetrameric complex. D) BS3 cross-linking coupled to mass spectrometry revealed extensive interactions between Ssu72 and Symplekin, primarily within Symplekin residues 1–320. Additional crosslinks were detected between WDR82, the Symplekin NTD, and a small region of PNUTS, in agreement with biochemical and structural analyses. E) WDR82 mediates interaction of PNUTS with CPA. Coomassie-stained SDS–PAGE and anti-FLAG immunoblotting indicates that PP1 associates with PNUTS independently of WDR82 (lanes 6–7), whereas efficient CPA binding required WDR82 (lane 7). No PP1 or CPA were detected in the PNUTS-free control (lane 8).

Symplekin, interacts with and activates the CPA-associated Pol II Ser5-P phosphatase Ssu72 in humans^78^ and Symplekin -Ssu72 can be reconstituted as a part of the Phosphatase module in *S. cerevisiae* CPF^93,94^. We therefore hypothesised that PPW interacts with 3’end processing machinery via Symplekin. To further elucidate how PPW interacts with CPF, we cloned PP1-PNUTS-WDR82-Symplekin-Ssu72 into a baculovirus construct and attempted to purify the heteropentameric complex using a twin-StrepII tag on Symplekin. While mass spectrometry confirmed the presence of all five subunits in the resulting complex, PP1 was markedly sub-stoichiometric, which we attribute to inefficient expression and folding in insect cells (Figure 3C). Nevertheless, our data indicate that Symplekin and Ssu72 can form a heterotetrametric complex with PNUTS and WDR82 within CPF. To probe interactions within this PNUTS-WDR82-Symplekin-Ssu72 complex, we employed amine-reactive bis(sulfosuccinimidyl) suberate (BS3) cross-linking coupled to mass spectrometry (XL-MS). We observed numerous crosslinks between Ssu72 and Symplekin, most of which were localised between residues 1 to 320 of Symplekin, consistent with reported structures of Symplekin-Ssu72 complexes from humans and *D. melanogaster*^78,95^ (Figure 3D). We also observed crosslinks between WDR82, Symplekin_NTD_, and a small region within PNUTS residues 565 and 591 (Figure 3D). We conclude that in addition to binding Ssu72, Symplekin_NTD_ directly interacts with WDR82 and PNUTS, and may link the PPW complex to the CPA machinery.

To test this hypothesis directly, we performed *in vitro* reconstitution experiments. We reconstituted a variant of *S. pombe* CPF lacking PNUTS, WDR82, and PP1, which we term core CPF, through co-expression of the remaining 10 subunits divided into two MultiBac constructs in Sf9 cells. Full-length Strep-tagged PNUTS was immobilised and incubated with PP1 and the core CPF in the presence or absence of WDR82. Following washes, the bead-bound fractions were analysed by Coomassie-stained SDS-PAGE and Western blotting with anti-FLAG antibody to allow unambiguous detection of FLAG-tagged PP1 (Figure 3E). Notably, while PP1 bound to PNUTS independently of WDR82 (Figure 3E, lanes 6 and 7), the CPA core associated with PNUTS and PP1 only in the presence of WDR82 (Figure 3E, lane 7) suggesting that WDR82 plays a central role in bridging the PPW complex and CPA complexes.

### Structural insight into PPW assembly with the 3’end processing machinery

We next sought to understand how WDR82 recruits PNUTS and PP1 to CPF via Symplekin. Our *in vitro* reconstitution and crosslinking data show that WDR82 directly interacts with PNUTS and the N-terminal domain of Symplekin. Furthermore, PNUTS and WDR82 form a stable heterodimer (Figure S3A). Our data support a model where WDR82 bridges PP1-PNUTS to the CPA. Therefore, we used our XL-MS data together with AlphaFold2 to define the WDR82-binding region of PNUTS (PNUTS_WBR_), which we predict to encompass PNUTS residues 565-644 (Figure 4A). We confirmed this prediction by expressing PNUTS_WBR_-WDR82 and determining its structure by X-ray crystallography to 1.6 Å resolution. All residues of PNUTS_WBR_ as well as all residues of WDR82 except for the final two were successfully modelled (Figure 4B). PNUTS_WBR_ interacts extensively with WDR82, burying a surface area of 9478 Å^2^ (Figure 4B). Most interactions between PNUTS_WBR_ and WDR82 are hydrophobic and ionic in nature. A stretch of hydrophobic residues _565_EIIWYKPVPIKF_576_ (WPL/I) makes contacts along a groove in WDR82, whilst another stretch of hydrophobic interactions is mediated by residues 639-641 of PNUTS, which we refer to as the LI -like motif (Figure 4B and C). Both the WPL/I and LI motifs are highly conserved (Figure 4C). The region between 586-590 (_586_RGYKC_590_) forms a half-helix over the centre of the WD40 domain of WDR82, inserting K_589_ into the central pore. Additionally, PNUTS _599_PEATSEIEREKN_610_ forms a helix enriched in charged residues providing another interaction site.

**Figure 4.**
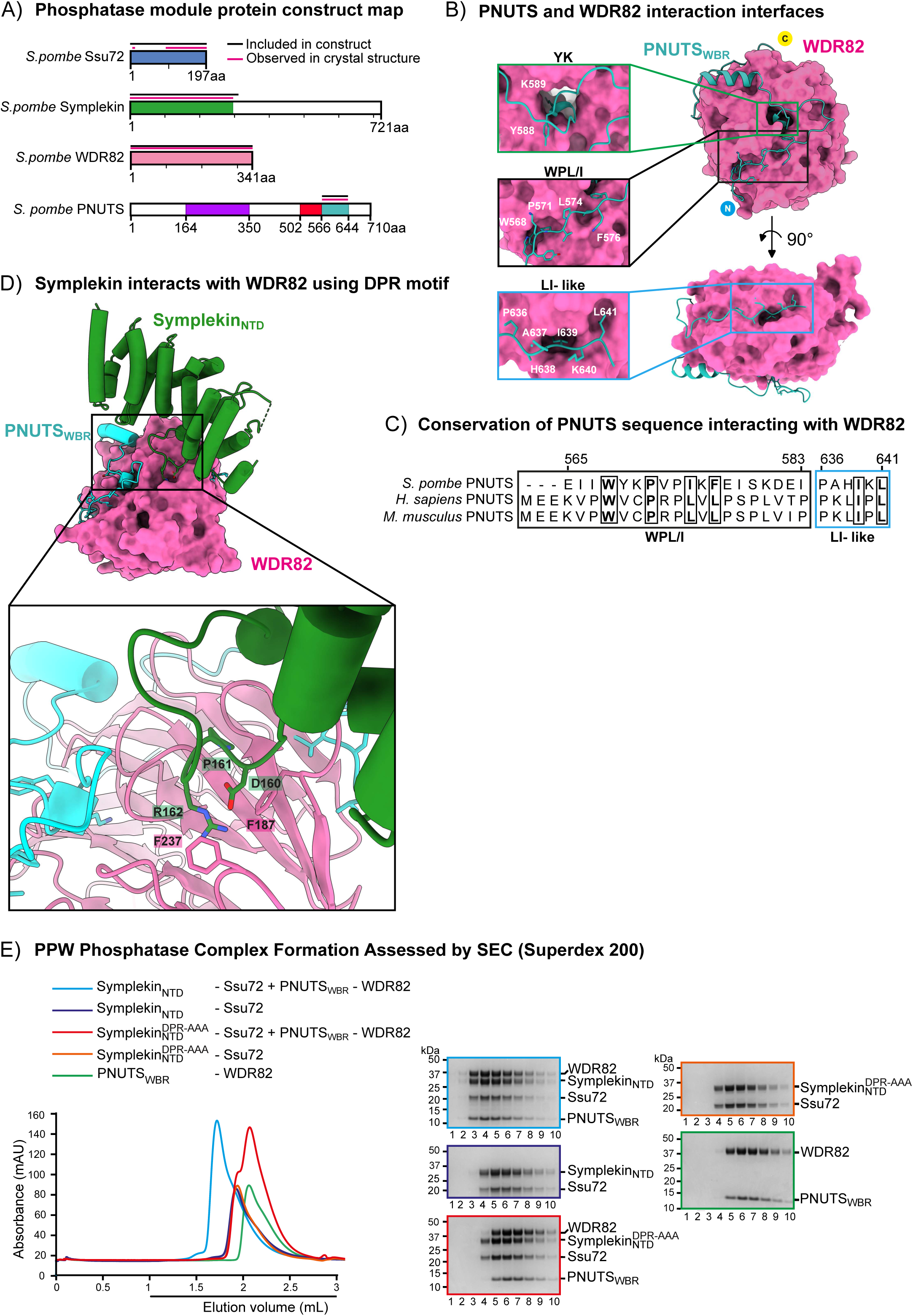
A) Domain maps of Symplekin, PNUTS, Ssu72 and WDR82 from *S. pombe.* Regions included in the X-ray crystallography constructs are indicated via black line, and the portions resolved in the crystal structure are highlighted via pink line. B) The PNUTS_WBR_-WDR82 complex structure was solved by X-ray crystallography. PNUTS_WBR_ makes extensive hydrophobic contacts with WDR82, mediated by conserved WPL and LI motifs, along with additional helix–groove interactions. C) Conservation of PNUTS -WDR82 interacting motifs (WPL/L and LI-like motifs). D) Microseeded crystallisation yielded a 3.8 Å structure of the reconstituted PNUTS_WBR_–WDR82–Symplekin_NTD_–Ssu72 complex. Symplekin_NTD_ forms a HEAT-repeat domain that contacts WDR82 through a conserved DPR motif in its extended α7–8 loop, and PNUTS provides additional stabilising contacts. E) Analyses of the complex for PNUTS_WBR_ –WDR82 with Symplekin_NTD_ –Ssu72 or Symplekin ^DPR-AAA^-Ssu72 formation by analytical SEC (Superdex200). Symplekin ^DPR-AAA^ mutant formed a heterodimer with Ssu72, however, failed to assemble with PNUTS_WBR_ –WDR82 into the hetero-tetramer, in contrast to wild-type SymplekinNTD –Ssu72.

We then reconstituted a heterotetrameric PNUTS_WBD_–WDR82-Symplekin_NTD_–Ssu72 complex by mixing individually purified PNUTS_WBD_–WDR82 and Symplekin_NTD_–Ssu72 complexes, followed by size-exclusion chromatography (Figure S3B), and crystallised the resulting complex, which diffracted to 3.8 Å resolution. We used the experimentally determined PNUTS_WBR_–WDR82 crystal structure, as well as AlphaFold2-generated models of Symplekin_NTD_ and Ssu72, as initial search models to solve the tetramer structure by molecular replacement. Interestingly, the density for Ssu72 is poor and is only visible for most of the low-molecular-weight protein tyrosine phosphatase (LMWPTP) fold, which constitutes the core of Ssu72. The density for the cap domain, as well as a helix and a β-strand from the LMWPTP was absent from the structure (Figure S3C, D). The portion of Ssu72 that could be modelled includes the interface with Symplekin_NTD_, with critical residues reported for the human Symplekin_NTD_–Ssu72 interface being highly conserved (Figure S3E). While we do not fully understand the reason for the missing electron density, we speculate that the Ssu72 cap domain is flexible relative to the Symplekin-bound LMWPTP domain and adopts multiple orientations within the crystal. Within the tetrameric complex, the PNUTS_WBR_–WDR82 heterodimer maintains the same overall conformation as observed when PNUTS_WBR_–WDR82 structure solved independently (Figure S3F).

Symplekin_NTD_ is composed of twelve anti-parallel helices arranged in a HEAT domain arrangement like its metazoan counterpart and contacts both Ssu72 (Figure S3E) and WDR82 (Figure 4D). The loop connecting α–helices 7 and 8 (hereafter referred to as the α7–8 loop) is exceptionally long, with the tip of the loop protruding outwards from Symplekin and inserting into a ‘pocket’ in WDR82 formed by two blades of the WD40 domain (Figure 4D). The tip of the α7–8 loop is formed by three amino acids - Asp_160_, Pro_161_, and Arg_162_, hereafter referred to as the DPR motif. Proline 161 is in a *cis* configuration and maintains the sidechains of the flanking Asp_160_ and Arg_162_ residues in an outward-facing orientation (Figure S4A). The sidechains of the flanking Asp_160_ and Arg_162_ make electrostatic interactions with each other, further stabilising the orientation of this DPR motif (Figure S4A). The aspartic acid (Asp_160_) and arginine (Arg_162_) residues of the DPR motif form ion–π interactions with Phe_187_ and Phe_237_ of WDR82 (Figure S4A and 4D).

Interestingly, the DPR motif was previously shown *in vivo* to be required for interaction of WDR82 with SET1A and ZC3H4 in mammals as part of the SET1 and Restrictor complexes, respectively^69,96^. We therefore hypothesized that the DPR motif is an evolutionary-conserved motif utilized by different WDR82-interactors. To test the importance of the Symplekin DPR motif observed in our crystal structure for binding WDR82, we expressed and purified a variant of the Symplekin _NTD_–Ssu72 heterodimer in which the DPR motif was replaced with three alanines. Symplekin ^DPR-AAA^ was fully functional in the formation of a heterodimer (Symplekin ^DPR-AAA^–Ssu72) with Ssu72 and eluted in the same volume as the Symplekin _NTD_–Ssu72 heterodimer in analytical size exclusion chromatography (Figure 4E, orange), showing that Symplekin binding to Ssu72 is unaffected by the loss of the DPR motif. However, while mixing Symplekin_NTD_–Ssu72 and PNUTS_WDB_–WDR82 resulted in the formation of the hetero-tetramer (Figure 4E, blue), Symplekin ^DPR-AAA^–Ssu72 did not interact with PNUTS_WBD_–WDR82 (Figure 4E, red). This demonstrates that the DPR motif in Symplekin is essential for binding WDR82. Our structure additionally shows limited interaction between PNUTS_WBR_ and Symplekin_NTD_, suggesting that Symplekin and PNUTS bind WDR82 independently. To test this, we repeated our reconstitution experiments without PNUTS and evaluated the results by analytical SEC. Our results showed that Symplekin_NTD_–Ssu72 bound WDR82, whereas Symplekin ^DPR-AAA^–Ssu72 did not, confirming that Symplekin and PNUTS bind WDR82 independently (Figure S4B). Moreover, these experiments suggest that PP1 and Ssu72 do not directly interact.

To further delineate the importance of different regions of PNUTS engaged in the interaction with WDR82, we generated PNUTS construct in which residues 566–591 within the WBR, comprising the highly conserved WPL/I (Figure 4C) and positive/negative helix were replaced with a glycine-alanine linker (referred to as PNUTS^WPL-G/A^) (Figure S4C, D). This mutant retained the ability to pull down PP1 but lost the ability to pulldown Symplekin _NTD_–Ssu72 even in the presence of WDR82 (Figures S4D, lanes 5-8). This data suggests that the WPL/I motif and the helix are essential for the PNUTS - WDR82 interaction, explaining its conservation (Figure 4C), and this interaction is required to link PNUTS–PP1 with Symplekin–Ssu72.

To dissect the role of the Symplekin DPR motif in mediating the CPA–PPW interaction, we introduced alanine substitutions at the DPR motif (DPR→AAA) to generate the Symplekin ^DPR-AAA^ mutant *in vivo*. Symplekin is an essential gene (Figure S5A), therefore we introduced an additional copy of the Symplekin gene with the DPR-AAA mutation into a strain in which the endogenous Symplekin gene was tagged with an Auxin-inducible degron (AID) (Figure 5A). AID-tagged Symplekin was efficiently depleted after growing cells for 1hr (Figure S5B) and remained depleted following 24 hours in the presence of 1 mM auxin. Complementing the Symplekin-AID strain with a wild-type copy of Symplekin verified that Symplekin^WT^ and Symplekin^DPR-AAA^ were expressed at similar levels (Figure 5B). Complementation with Symplekin^WT^ but not Symplekin^DPR-AAA^ rescued growth after auxin-induced depletion of endogenous Symplekin (Figure 5C), suggesting that the interaction between CPA and PPW mediated by the DPR motif in Symplekin is essential for cell viability.

**Figure 5.**
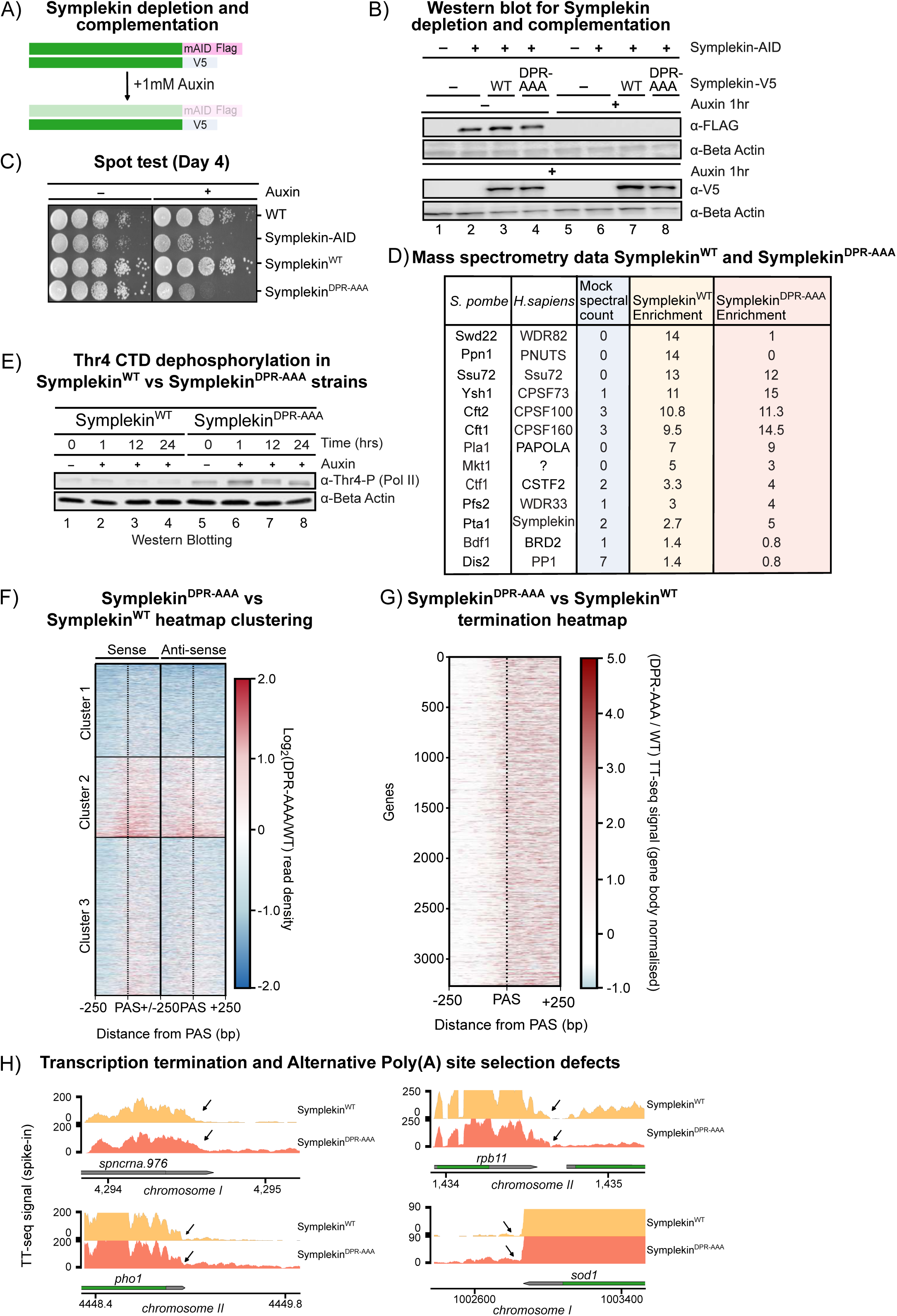
A) Schematic of endogenous Symplekin with AID and FLAG-tag, and secondary copies of Symplekin^WT^ and Symplekin^DPR-AAA^ with V5-tag. B) Western blot analyses of Symplekin^WT^ and Symplekin^DPR-AAA^ with β-actin used as a loading control. C) Spot test on YES Auxin plates of WT, Symplekin-AID, Symplekin^DPA-AAA^ and Symplekin^WT^. Symplekin^DPA-AAA^ shows a large growth defect not observed in the Symplekin^WT^ strain. D) Purification of V5-tagged Symplekin^WT^ and Symplekin^DPR-AAA^ followed by proteomic analysis of interacting partners, using mock without V5 tag as control. E) Pol II Thr4-P levels were assessed after depletion of AID–Symplekin in strains complemented with Symplekin^WT^ or Symplekin^DPR-AAA^. The Symplekin^DPR-AAA^ mutant elevated Thr4-P even before depletion, and further increases were observed upon loss of endogenous Symplekin. F) TT-seq analysis of Symplekin^WT^ and Symplekin^DPR-AAA^ following auxin treatment. Representative heatmap of gene expression +/- 250 base pairs centred around the PAS, presented as log_2_ fold ratio relative to Symplekin^WT^. Sense and antisense strands presented as log_2_ fold ratio relative to Symplekin^WT^. G) Heatmap representing termination defects (readthrough), scaled by expression in the gene body presented as log_2_ fold ratio relative to Symplekin^WT^. H) Representative snapshots of Symplekin^WT^ and Symplekin^DPR-AAA^ expression. Arrows highlight proximal PAS used predominantly in WT cells.

Analysis of native purifications of V5-tagged Symplekin by mass spectrometry revealed that, in contrast to WT Symplekin, which co-purifies with the entire CPF complex including PPW, Symplekin^DPR-AAA^ lost the ability to interact with PPW. However, Symplekin^DPR-AAA^ retained the ability to interact with the rest of CPF, demonstrating that the overall integrity of the CPA is not compromised by the DPR-AAA mutation (Figure 5D). PP1 interacted nonspecifically with V5 beads, as evidenced by its presence in the no-tag control (mock). These results therefore confirm our conclusions from *in vitro* experiments that the DPR motif of Symplekin is important for interacting with WDR82 and recruitment of PNUTS, WDR82, and PP1 to the CPA machinery.

To determine if the interaction between PPW and the CPA is necessary for dephosphorylation of PP1 substrates involved in transcription, we assessed phosphorylation of Thr4 of the Pol II CTD upon depletion of the endogenous Symplekin in cells complemented with Symplekin^WT^ or Symplekin^DPR-AAA^. Interestingly, expression of Symplekin^DPR-AAA^ before depletion of AID-tagged Symplekin^WT^ accumulates hyper-phosphorylated Thr4-P Pol II, suggesting that Symplekin^DPR-AAA^ may have a dominant negative effect (Figure 5E, compare lanes 5 to 1). Depletion of endogenous AID-tagged Symplekin leads to a further increase in the levels of phosphorylated Pol II (Figure 5E, lane 6), demonstrating that association of the PPW complex with the CPA machinery is important for dephosphorylation of Pol II by PP1.

Collectively, our data demonstrate that WDR82 plays a critical role in bridging PNUTS-PP1 to the CPA machinery via Symplekin. Both PNUTS and WDR82 are important for dephosphorylation of PP1 substrates *in vivo*, but only PNUTS can directly stimulate PP1 activity. In contrast, our structural and biochemical data demonstrates that WDR82 plays a prominent role in mediating interaction between CPA and PPW complexes by interacting with the DPR motif of Symplekin. PNUTS on the other hand, is important for mediating interaction between WDR82 and PP1.

### Loss of physical link between CPA and PP1 phosphatase complex leads to altered 3’end processing and transcription termination defect

We next investigated the global transcriptional consequences of disrupting the interaction between the CPA complex and the PPW complex. To this end, we performed spike-in normalised transient transcriptome sequencing (TT-seq) in the Symplekin^WT^ or Symplekin^DPR-AAA^ cells. Analysis of nascent transcription revealed a widespread increase in read density downstream of annotated PAS in the Symplekin^DPR-AAA^ strain compared to Symplekin^WT^. The readthrough past the annotated TES was particularly pronounced for genes in cluster 2 (Figure 5F), and a mild reduction in TT-seq signal across gene bodies was also observed. The reduction in signal across gene bodies does not correlate strongly with antisense transcription (Figure 5F). Additionally, we normalised the post-PAS signal to expression levels to accurately quantify the termination defects. Normalisation confirmed that the 3′-end processing and termination defects associated with Symplekin^DPR-AAA^ are globally affecting transcription across the genome (Figure 5G). For a subset of genes, readthrough also reduced transcription of the neighbouring gene (tandem orientation of the genes). For example, at the *rbp11–spac3a12.08* locus, readthrough from *rbp11* (encoding a Pol II subunit) interferes with transcription of the downstream *spac3a12.08* gene (encoding an acyl-CoA thioesterase) revealing how readthrough transcription could negatively impact gene expression from the adjacent promoter (Figure 5H). We next compared the readthrough defects in Symplekin^DPR-AAA^ against the readthrough defects in the PPW mutants. Genes with increased levels of 3’readthrough, observed in at least two PPW mutants, showed a substantial overlap with genes exhibiting readthrough in the Symplekin^DPR-AAA^ strain (Figure S5C, left). This trend was consistently observed across multiple comparisons (Figure 5C, right).

We further examined alternative polyadenylation (APA), using the same pipeline applied to the PPW mutants (Figure 2D, S2C–D). Symplekin^DPR-AAA^ complementation resulted in alternative polyA site selection in a subset of genes; these alterations did not correlate with gene length and showed only minor differences in GC content relative to unaffected genes (Figure S5D–E). Analysis of non-coding RNAs in the Symplekin^DPR-AAA^ complemented strain revealed no significant alterations, although a slight decrease in rRNA levels was observed compared with Symplekin^WT^ (Figure S5F). In summary, we identified a direct functional link between CPA and the PPW complex, and our findings demonstrate that coordinated phosphatase activity and CPA are integral to proper 3′-end processing and transcription termination. Our data supports a model in which PP1, activated by PNUTS, is recruited to elongating Pol II via WDR82 and Symplekin to coordinate Spt5/CTD dephosphorylation with pre-mRNA 3′-end cleavage and polyadenylation (Figure 6A).

**Figure 6.**
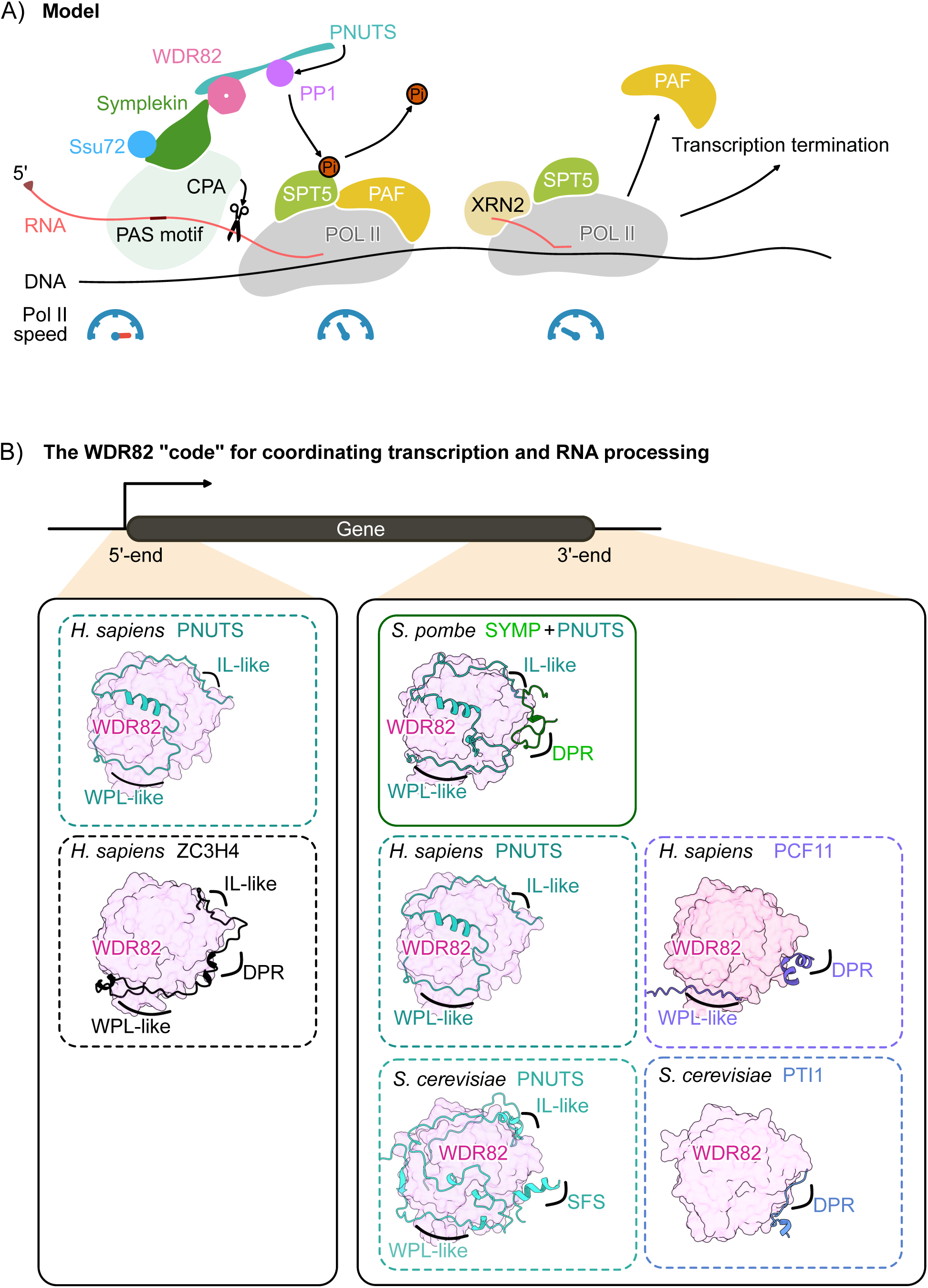
A) Model describing how PAS recognition by the CPA is linked to dephosphorylation of Spt5 via recruitment and activation of PP1 by WDR82 and PNUTS. Symplekin acts as a scaffold that links the phosphatase holoenzyme (PPW) complex to the 3’end processing CPA machinery via the DPR motif. WDR82 bridges this interaction, while PNUTS not only recruits PP1 but also enhances its activity toward Spt5. Dephosphorylation of Spt5 can subsequently slow Pol II, promote dissociation of the PAF complex, and facilitate 3′-end processing and transcription termination. B) WDR82 functions as a central hub that coordinates multiple machineries involved in transcriptional regulation at both the 5′ and 3′ ends of genes. Building upon the structural data presented in this study (solid box), we employed AlphaFold modelling (dashed boxes), to predict how conserved SLiMs could engage with WDR82 to mediate formation of the diverse set of complexes.

## Discussion

Transcription termination requires the action of phosphatases to remove phosphorylation marks associated with productive elongation from RNA polymerase II and associated transcription factors^97^. The PPW holoenzyme has emerged as a conserved central regulator of the transition from transcription elongation to termination, acting at the 3′ ends of genes and premature termination sites across eukaryotes, from yeast to humans^43–46,49–52,69,84–87,96,98–102^. Here, we define the mechanism by which the PPW holoenzyme mediates transcription termination by uncovering its role in pre-mRNA 3′-end processing, and by elucidating how PP1 is recruited and regulated in fission yeast. Our findings further suggest that analogous interaction networks operate broadly across eukaryotes.

Our data supports a model in which PP1, activated by PNUTS, is recruited to elongating Pol II via WDR82 and Symplekin, thereby coordinating Spt5/Pol II dephosphorylation with pre-mRNA 3′-end cleavage and polyadenylation (Figure 6A). In this framework, WDR82 acts as a hub that recognises multiple short linear motifs (SLiMS) in PNUTS and Symplekin via conserved residues (Figures 4B-D and S6A), creating a platform where the PP1 holoenzyme, the CPA machinery, and Pol II converge at the 3′ ends of genes.

By defining the modular interfaces connecting PP1, PNUTS, WDR82, and Symplekin, this work reveals how the conserved PPW complex couples transcription with RNA 3′-end processing to promote dephosphorylation of the transcription machinery during the elongation–termination transition. Our results revise the prevailing view that PP1 functions primarily downstream of RNA cleavage and instead support a model in which PPW recruitment to the CPA modulates PAS selection and transcription termination We propose that the PPW complex coordinates PAS selection and cleavage efficiency through several potential mechanisms. One possibility is that PPW association with the CPA induces conformational changes within the 3’end processing machinery, thereby influencing PAS recognition or modulating the efficiency of endonucleolytic cleavage. This could be either caused by the PPW-mediated dephosphorylation of CPA subunits, which may alter its affinity or specificity for PAS elements and/or influence the pre-mRNA cleavage efficiency by CPSF73/Ysh1. It is also possible that PNUTS could facilitate recruitment of CPA to PAS by interacting with nascent RNA via its zinc finger domain, which is present in mammalian PNUTS (Figure 1A)^106^. Secondly, we propose that PPW facilitates dissociation of elongation factors from Pol II, inducing rearrangements in the elongation complex and slowing Pol II elongation dynamics. PP1-mediated dephosphorylation of Spt5 was reported to slow Pol II down near gene ends, and we propose that Pol II deceleration extends the time window for CPA recognition of the PAS and promotes efficient cleavage. Dissociation of the elongation factors could make Pol II more susceptible to dislodgement by Xrn2 due to allosteric changes in the Pol II-DNA-RNA complex leading to its destabilization, which is consistent with a unified allosteric-torpedo model of termination^103^.

Notably, several lines of evidence indicate that PPW promotes transcription termination primarily by remodelling the elongation complex and slowing Pol II, rather than by directly regulating RNA cleavage. PP1, PNUTS, and WDR82 can mediate cleavage-independent termination via the NNS complex in budding yeast and the Restrictor complex in metazoans ^69,72,74,75,79–81,85,86,104,105^, demonstrating that the PPW complex can drive transcription termination independently of CPA endonucleolytic activity. Consistently, the Restrictor has been proposed to drive transcription termination by slowing Pol II speed^86^. Additionally, PP1-mediated transcription termination requires dissociation of the PAF complex^17^, which can lead to rearrangements of the Pol II elongation complex and a reduction in Pol II speed^107,108^. Finally, dephosphorylation of Spt5 by PP1 is associated with deceleration of Pol II at the 3’ end of genes, and Pol II speed has been proposed to contribute to the recruitment of RNA processing factors ^109–118^.

We present the first experimental structure of a PNUTS-WDR82-Symplekin-Ssu72 complex, demonstrating that PNUTS and Symplekin bind to WDR82 via distinct interfaces, each using well-defined SLiMs. We define three conserved WDR82-interacting SLiMs: WPL/I, LI-like, and DPR motifs. In our experimental structures, PNUTS utilises WPL/I and LI-like motifs while Symplekin uses a DPR motif to form the PPW complex. We demonstrate that the physical link between PPW and CPA is important for transcription termination and pre-mRNA 3’end processing. Comparison of Symplekin homologs from yeast to humans showed that the DPR motif is present only in fission yeast Symplekin (Figure S6A-D).

While PNUTS, WDR82, and PP1 homologs have now been identified across a range of eukaryotes, only in yeasts are these proteins stably integrated into the CPA. Metazoan Symplekin lacks a DPR motif, which explains why mammalian PNUTS-WDR82-PP1 does not stably co-purify with the CPSF components ^18,119^. Despite the absence of the DPR motif in human Symplekin, purification of human 3’end processing complexes using PAS-containing RNA as a bait, revealed the presence of PNUTS and PP1 (but not WDR82) in addition to the CPA components^119^. Reciprocally, the CPA was shown to co-purify with mammalian PNUTS, together with WDR82 and PP1^51^. Together, these data suggest that a functional link between PPW and the CPA is conserved across eukaryotes, albeit potentially mediated through different CPA subunits or alternative interaction surfaces. Interestingly, WDR82 has been proposed to interact with the CFIIm component hPCF11 to promote premature transcription termination during HIV infection^98^. The WDR82–hPCF11 interface has not been defined experimentally; however, hPCF11 contains both a DPR motif and a WPL-like motif, analogous to those found in PNUTS (Figure 6B). AlphaFold modelling suggests that these motifs engage the same DPR pocket and WPL groove on WDR82 as observed in the PPW–Symplekin structure (Figures 6B and 4B, C, D). Therefore, although the position of the DPR motif is not conserved across partners, the underlying binding principles are maintained. Additionally, the absence of a DPR-motif in budding yeast Symplekin is surprising given that orthologs of PNUTS, WDR82, and PP1 exist as a stable submodule of CPF together with other *S. cerevisiae* specific subunits – Syc1 and Pti1^93,94^. Due to the highly conserved nature of the DPR-binding pocket, we speculate that the DPR motif resides in another CPA subunit in *S. cerevisiae.* Interestingly, the budding yeast ortholog of PNUTS (Ref2) has WPL/IL and DPR motifs (Figure 6B). However, the CPF subunit Pti1 contains a DPR motif and is also part of the CPF phosphatase module, Pti1 could potentially be the interaction partner of *S. cerevisiae* WDR82 (Figure 6B).

Consequently, we propose that while the subunits contributing to the PPW–CPA interface have diverged between species, the overarching principle of PPW engagement with the 3′-end processing machinery/ termination complexes is evolutionarily maintained. Crucially, the binding grooves of these SLiMs on WDR82 are distinct and non-overlapping, suggesting that multiple WDR82 interactors can co-occupy the same molecule of WDR82, opening the possibility of a combinatorial WDR82-binding code. An open question is what the molecular rules governing this code are, especially in cases where a protein contains WPL/I and LI-like motifs, as well as a DPR motif. This is especially pertinent to metazoan WDR82, which is an integral subunit of multiple complexes that regulate transcription in distinct and even antagonistic ways. In addition to the PPW complex, WDR82 also binds ZC3H4 to form the Restrictor complex, which restricts non-coding transcription in metazoans, through a DPR motif in the ZC3H4 subunit^69,85^. Interestingly, Restrictor was originally proposed to consist solely of ZC3H4 and WDR82, and it was suggested that PNUTS and ZC3H4 binding to WDR82 would be mutually exclusive. Indeed, AlphaFold predictions of the binary ZC3H4-WDR82 complex show that ZC3H4 forms extensive interactions with WDR82, in addition to the DPR-binding pocket, through the WPL/IL binding grooves of WDR82 (Figure 6B).

However, our data does not rule out the possibility that hybrid complexes could be formed via different Short Linear WDR-Interacting motifs (WDR-SLiMs). Indeed, AlphaFold predictions of a trimeric PNUTS-WDR82-ZC3H4 complex show PNUTS occupying the WPL/I and LI-like grooves while ZC3H4 occupies only the DPR-pocket, in remarkable agreement with our experimental PNUTS-WDR82-Symplekin-Ssu72 structure (Figure S6E, F). In support of this, ZC3H4 has recently been shown to copurify with the PPW complex^85^. WDR82 is also a part of the H3K4 histone methyltransferase SET1 complex, in *S. cerevisiae* and in mammalian cells^96,101,120,121^ . WDR82 binding to the SET1A subunit of the H3K4 histone methyltransferase SET1 complex has been proposed to antagonise Restrictor function in certain loci, through a DPR motif in SET1A. Yet, PNUTS has not been identified in SET1 complexes. SET1A would also occupy the groove in WDR82 occupied by PNUTS via WPL/IL motifs, however, with opposite polarity, similar to ZC3H4 (engaging motif ‘LxxPxF’ for SET1 and ZC3H4 as opposed to ‘WxxPxxL/I’ in PNUTS as revealed by our PNUTS_WBR_ – WDR82 structure through hydrophobic interactions, which are conserved. Incorporation of WDR82 into the SET1 and 3’end processing/termination complexes has been proposed to be mutually exclusive, as reported for Restrictor-ZC3H4 in mammalian cells^69^ . This may apply to the NNS complex, as WDR82 was shown to be important for APT recruitment to snoRNA genes that rely on NNS for termination^72,75,101^ However, a physical link between NNS and the APT complex has not yet been reported. These data support the prediction that SET1A indeed occupies all binding surfaces on WDR82 (Figure S6G). These seemingly contradictory observations underscore the need to validate *in silico*-predicted structural models experimentally. Future structural studies with experimental techniques will be required to validate this combinatorial WDR82-binding code.

This principle could have broader implications beyond the recruitment of the PPW complex to RNA processing machineries. Interestingly, WDR82 was proposed to interact with and mediate the recruitment of the silencing HUSH complex to sites of transcription termination^122^. The HUSH complex has been implicated in the suppression of foreign DNA expression by mediating the recruitment of chromatin remodelling activities and the H3K9 histone methyltransferase. Interestingly, TASOR subunit of HUSH has a DPR motif and WPL-like motifs that could mediate the interaction with WDR82. Additionally, this principle is not limited to WDR82 and is likely to apply to other WD40 repeat proteins. For example, the WD40 repeat factor CstF-50, which is part of the mammalian Cleavage Stimulation factor (CstF), partly interacts with CstF77 via a motif similar to DPR (DPK) to aid recognition of the G/U-rich elements downstream of PAS and PAS selection by the CPA machinery and cleavage factors^123^. Similar to WDR82, CstF-50 might mediate links to other complexes and was proposed to interact with the tumour suppressor E3-ubiquitin ligase complex BRCA1 (subunit BARD1)^124^.

We propose that WDR82 recognises common SLiMs across its interactors, with distinct WDR-interacting motifs enabling a combinatorial interaction code. This combinatorial WDR82-binding code dynamically regulates phosphatase activity, RNA processing, chromatin modification, and transcription termination. Analogous to a molecular “Lego” system, this modular interaction system is conserved from yeast to humans. By revealing this code, our work establishes WDR82 as a decisive integrator of transcriptional control and uncovers a general principle by which WD40 repeat proteins organise and couple fundamental cellular processes.

## METHODS

### Strain construction and growth

*S. pombe* strains were grown in YES medium at 30°C to OD_600_ of 0.4 - 0.7, otherwise indicated. Standard homology-based methodology for genomic integration was used for either epitope tagging or gene deletion. Tag integration was validated by Western Blotting with antibodies against α-FLAG (A8592, Sigma-Aldrich), α-V5 (V226, Sigma-Aldrich or MCA1360, Bio-Rad), and α-tubulin (ab6160, Abcam). Complementary copy of V5-tagged Symplekin either WT or DPR to AAA mutant was targeted to *ura4* locus under native promoter to complement for the depletion of the endogenous mAID-FLAG-tagged Symplekin/Pta1. Depletion of mAID-FLAG-tagged Symplekin were performed in YES in the presence of 1 mM auxin (Sigma) at 25°C. Strains generated and used in this study are listed in Supplementary Table 1.

### Immunofluorescence

Immunofluorescence of PNUTS-FTP was carried as published^32,43^. Briefly, 9 ml of yeast culture (OD_600_ = 0.5) were fixed by addition of 1 ml 37% formaldehyde for 30 min at RT with mixing on a rotating wheel. Cells were washed three times with PEM (100 mM piperazine-N,N′-bis[2-ethanesulfonic acid] /PIPES/, 1 mM EGTA pH 8.0, 1 mM MgSO_4_, pH 6.9) and treated with 12.5 µl 10 mg/ml zymolyase T100 (9329.2, Carl Roth) in 500 µl PEMS (PEM with 1.2 M sorbitol) for 1h at 37°C. Cells were permeabilised in 500 µl PEMS, 1% Triton X-100 for 30 s, washed three times with PEM, then incubated on a rotating wheel for 30 min in 200 µl PEMBAL (PEM with 1% bovine serum albumin, 100 mM lysine hydrochloride, 0.1% NaN_3_) at RT. 1/5^th^ of total fixed cells were used per condition. Cells were incubated with α-FLAG (Sigma, F3165, 1:200) in 100 µl PEMBAL at 4°C overnight, then washed twice with PEMBAL, followed by incubation in 200 µl PEMBAL for 30 min at RT. Cells were stained with Alexa Fluor 488-conjugated Goat α-Mouse IgG (H+L) Cross-Adsorbed Secondary Antibody (Invitrogen, A11001, 1:400) in 100 µl PEMBAL at 4°C overnight in the dark. Cells were washed once with PEMBAL and once with PBS, then resuspended in 40 µl PBS. Next, 3 µl cell suspension were freshly mounted on poly-lysine-coated coverslips, let to dry, and imaged under ROTIMount FluorCare DAPI (HP20.1, Carl Roth). Z-stacks were acquired on a Deltavision Ultra High-Resolution Microscope (Cytiva) using AcquireUltra (1.2.2) and deconvolved with softWoRx (7.2.1) using default settings. Image quantification was carried out with Fiji (ImageJ 1.53t) using a semiautomated ImageJ macro script (FISHquant_v6.ijm; https://github.com/Kilchertlab/Microscopy-analysis). In short, cellular segmentation was carried out by thresholding the transmitted light channel (cell outlines) and DAPI stain (nuclei). FLAG signal was measured on average intensity Z-projections for nucleus and cytoplasm (total cell without nucleus) and the ratio of mean nuclear fluorescence intensity over mean cytoplasmic fluorescence intensity calculated for each cell.

### Purification of native PP1-3xFLAG from *S. pombe*

Strain with the C-terminally FLAG-tagged PP1 was grown in 48 L of 2X YES media (10 g/L Yeast Extract, 450 mg/L of Adenine, Histidine, Leucine, Uracil, Lysine hydrochloride, 6 g/L Glucose) at 30 °C. Cells were harvested when OD_600_ reached 5 by centrifugation using a JLA 8.1 rotor at 4000 g for 10 minutes at 4 °C, washed with PBS, and stored at -80°C. Cells were resuspended in Lysis Buffer (150 mM NaCl, 50 mM Tris pH 7.9, 10% glycerol, 0.5 mM MgCl_2_, 0.25% Triton X-100) supplemented with SuperNuclease (SinoBiological), EDTA-free Protease Inhibitor tablets (Roche) and lysed by French Press operated at 35 kPSI. Lysate was then supplemented with 0.1% Triton and 1 mM PMSF (phenylmethylsulfonyl fluoride) followed by centrifugation in a JLA 16.25 rotor for 30 minutes at 34000 g at 4 °C. The cleared lysate was incubated for 1 hour with 2.5 mL M2 Anti-FLAG resin (Sigma-aldrich) at 4 °C. Resin was then washed 3 times with 10 column volumes (CVs) of Lysis Buffer, followed by 3 times 10 CVs of Wash Buffer (500 mM NaCl, 50 mM Tris pH 7.9, 10% glycerol, 0.5 mM MgCl_2_, 0.5% Triton X-100, 500 mM Urea) , and 3 times with 10 CVs Lysis Buffer before elution using 5 mL 1 mg/mL FLAG peptide in Lysis Buffer. Eluate was diluted with QDA (20 mM NaCl, 50 mM Tris, 10% glycerol, 0.5 mM MgCl_2_, 1 mM β-mercaptoethanol) buffer to 50 mL and loaded onto a 5 mL HiTrap Q HP column (Cytiva) equilibrated with QDA buffer. Protein was eluted using a gradient of 0-100% QDB buffer (1 M NaCl, 50 mM Tris, 10% glycerol, 0.5 mM MgCl_2_, 1 mM β-mercaptoethanol) over 40 CV. Fractions containing PP1-3xFLAG were pooled, concentrated, snap-frozen in liquid nitrogen and stored at -80 °C.

### Purification of native PNUTS and Symplekin

FTP-tagged PNUTS and V5 tagged Symplekin were purified from yeast cells. As a mock control strain without tag was used. Cells were grown to OD 1.5, harvested and cell pellets were frozen at -80 °C. Cells were disrupted in a freezer mill (SPEX SamplePrep) in liquid nitrogen. For purification, 5 g of cell powder was resuspended in 25 mL of lysis buffer (buffer L) (25 mM Tris,pH 7, 100 mM NaCl, 0.5 mM MgCl_2_, 0.5 mM β-mercaptoethanol) supplemented with 1 mM PMSF, cOmplete Protease Inhibitor Cocktail (Roche) and phosphatase inhibitors: 1 mM NaF, 1 mM sodium orthovanadate, 2 mM imidazole, 1 mM sodium pyrophosphate decahydrate, 0.5 mM glycerol-phosphate, 5 µM cantharidin, 5 nM calyculin A. The lysates were incubated for 30 min in the cold room and cleared at 40000 x g for 20 min at 4 °C. Following centrifugation, the lysates were incubated with 250 µL of an equilibrated V5-Trap® Magnetic Particles M-270 (ChromoTek) slurry for 1 h in a cold room. Beads were washed 10 times with 1 mL of buffer L and proteins were eluted twice with 150 µL of 0.2 M glycine pH 2.5. Samples were neutralised with 25 µL of 1 M Tris. For tandem affinity purification, lysate was incubated for 2 hr at 4 C with 1 mL of IgG Sepharose beads (GE Heathcare). The beads were washed four times with 20 mL of lysis buffer then resuspended using 1 mL of lysis buffer. Proteins were eluted by incubation with 65 μl of TEV protease (0.5 mg) for 1 hr 30 min at 16 C. The sample was collected from the beads by centrifugation and incubated with 200 μL of M2 resin anti-FLAG (Sigma) beads for 1 hr at 4 °C. Beads were washed three times with 5 mL of buffer and the samples eluted with 100 μL glycine buffer and mixture was neutralised with 20 μL 1M TRIS. The samples were denatured with 8 M urea, reduced with TCEP, alkylated with 2-chloroacetamide, and digested with LysC, and trypsin. The reactions were stopped with formic acid and subjected to analysis by mass spectrometry.

### Recombinant protein expression and purification from *E. coli*

#### Recombinant PP1

A codon optimised *S. pombe* PP1/Dis2 (1–300 aa) coding sequence was cloned into pTXB1 to generate an N-terminal His₆ tag and a C-terminal Mxe GyrA intein–CBD fusion. The plasmid was transformed into *E. coli* BL21 cells co-expressing GroEL/GroES (pGRO7). Cultures were grown in 2xTY medium supplemented with chloramphenicol (30 µg/mL), ampicillin (50 µg/mL), and MnCl₂ (2 mM) at 37 °C. At OD_600_ = 0.5, GroEL/GroES expression was induced with 1 g/L L-arabinose and the temperature was reduced to 10 °C. When cells reach OD_600_= 1.0, PP1 expression was induced with 0.1 mM IPTG for 48 h at 10 °C. Cells were harvested, resuspended in fresh 2xTY medium containing MnCl₂ and chloramphenicol, incubated for 2 h at 10 °C, and stored at −70 °C. Pellets were lysed by sonication in lysis buffer (25 mM HEPES pH 7.9, 500 mM NaCl, 10% glycerol, 1 mM MnCl₂, 5 mM imidazole) supplemented with PMSF and EDTA-free protease inhibitors. Clarified lysates (40,000 g, 20 min, 4 °C) were applied to Talon Superflow resin (Takara Bio). Following, resin was washed and protein eluted with elution buffer (25 mM HEPES pH 7.9, 500 mM NaCl, 10% glycerol, 1 mM MnCl_2_, 250 mM imidazole), after which Intein self-cleavage was induced by a 10-fold dilution in cleavage buffer (25 mM HEPES pH 7.9, 500 mM NaCl, 10% glycerol, 1 mM MnCl_2_, 50 mM β-mercaptoethanol) and incubation at RT for 72 hours at 22 °C. The sample was diluted in lysis buffer and re-applied to Talon resin to remove uncleaved material. Eluted PP1 was dialyzed overnight at 4 °C into storage buffer (25 mM HEPES pH 7.9, 150 mM NaCl, 10% glycerol, 1 mM MnCl₂, 1 mM β-mercaptoethanol), snap-frozen in 40% glycerol, and stored at −70 °C.

#### PNUTS_420-710_ and PNUTS_420-710_ mutant variants

A gene fragment corresponding to *S. pombe* PNUTS residues 420 to 710 containing an N-terminal His₆ tag and a C-terminal Twin-Strep tag was cloned into a modified pTXB1 vector lacking intein-CBD. Mutants PNUTS^W510A^₄₂₀–₇₁₀ and PNUTS^R586/Y588/K589A^₄₂₀–₇₁₀ were generated by PCR mutagenesis. Proteins were expressed in *E. coli* Rosetta (DE3) cells grown in 2xTY medium supplemented with chloramphenicol (25 µg/mL) and ampicillin (50 µg/mL) at 37 °C. Upon reaching an OD_600_ of 1.0, protein expression was induced with 0.3 mM IPTG, followed by incubation at 20 °C overnight. Cell pellets were lysed in Strep buffer (50 mM HEPES pH 7.9, 150 mM NaCl, 0.5 mM MgCl₂, 1 mM β-mercaptoethanol) supplemented with PMSF and EDTA-free protease inhibitors. Lysates were clarified by centrifugation (40,000 g, 20 min, 4 °C) and filtration (0.22 µm) and applied to a 5 mL Strep-Trap HP column (Cytiva). Bound protein was eluted with Strep buffer containing 5 mM desthiobiotin (IBA Lifesciences) and subsequently incubated with Talon Superflow resin. After incubation, the resin was washed sequentially with Strep buffer containing 5 mM Mg²⁺-ATP and with 2 mM imidazole. Proteins were eluted using 250 mM imidazole. Final purification was performed by size-exclusion chromatography (HiLoad 16/600 Superdex 75 pg, Cytiva) equilibrated in gel-filtration buffer (25 mM HEPES pH 7.9, 500 mM NaCl, 10% glycerol, 1 mM MgCl₂, 1 mM β-mercaptoethanol). Peak fractions were concentrated using 10 kDa MWCO centrifugal filters, snap-frozen, and stored at −70 °C.

#### Spt5-Kow_5_-CTR_7_ and its phosphorylated form

N-terminally His tagged Spt5 KOW_5_-CTR_7_ was expressed in *E. coli* as described before^32^. Briefly, cleared lysate was affinity purified over Ni-NTA resin (Qiagen), followed application of protein to Heparin Sepharose (HiTrap Heparin HP, Cytiva) and polished on HiLoad 16/600 Superdex 75 (Cytiva). The Spt5-Kow5-CTR_7_ (54 μM) was phosphorylated with Cdk9_1-350_-Cyclin T (Pch1) kinase complex (0.4 μM) using 1 mM MgCl_2_, ATP and MnCl_2_ in a total volume of 1 mL STREP buffer (50 mM HEPES pH 7.9, 150 mM NaCl, 0.5 mM MgCl_2_, and 1 mM β-mercaptoethanol). The reaction was kept for 1.5 hours at 30 °C and then subjected to size exclusion chromatography using a HiLoad 16/600 Superdex 75 column (Cytiva). Peak fractions were combined and concentrated using an Amicon 10 kDa Ultra Centrifugal Filter (Millipore) and stored at -70 °C. The phosphorylation status of the sample was evaluated by changes in migration in a 10% acrylamide Phos-tag resolving gel (2.5 µM Phos-tag Acrylamide and 50 µM MnCl_2_).

### Recombinant protein expression from *Sf9* cells

All proteins expressed from Sf9 were cloned into pACEBac1. Bacmids were generated by transformation of DH10EmBacY cells and positive clones were screened by blue-white screening. Bacmids were extracted from 3 mL overnight cultures using isopropanol precipitation. 2 mL of Sf9 cells at 0.5 × 10^6^ were transfected with 2 μg of bacmid using FuGene transfection reagent (Promega). After 5 days at 27 °C, cells were harvested and the supernatant was filtered using a 0.22 μm filter. P2 virus was generated by infection of Sf9 cells at 1.5-2.0 × 10^6^ with P1 virus at a 1:100 ratio (v/v). After 4 days at 27°C, cells were harvested and supernatant was filtered using a 0.22 μm filter and stored at 4 °C in the dark. Large scale protein expression was carried out by infecting Sf9 cells at 2.0 × 10^6^ with P2 virus at a 1:100 ratio (v/v). Cells were harvested after 3 days at 27 °C, washed with PBS, and stored at -80 °C.

PNUTS, PNUTS^WPL-G/A^ and PNUTS-WDR82 were purified with the following protocol. Cells were resuspended in Strep Buffer (50 mM HEPES pH 7.9, 150 mM NaCl, 0.5 mM MgCl_2_, 1 mM β-mercaptoethanol) supplemented with SuperNuclease, EDTA-free Protease Inhibitor tablets (Roche), and 2 mM PMSF, and lysed by sonication. The lysate was clarified by centrifugation using a JA25.5 rotor at 40,000 g for 20 minutes at 4°C. The clarified lysate was filtered through a 0.22 μm filter and loaded onto a 5 mL StrepTrap HP column equilibrated with Strep Buffer. The column was washed with 20 column volumes of Strep buffer before elution with Strep Elution Buffer (Strep Buffer supplemented with 5 mM Desthiobiotin (IBA sciences)). The eluate was diluted to 30 mM NaCl and loaded onto a 5 mL HiTrap Q HP equilibrated with QA buffer (50 mM HEPES pH 7.9, 30 mM NaCl, 0.5 mM MgCl_2_, 1 mM β-mercaptoethanol). Protein was eluted with a gradient of 0 – 100% QB buffer (50 mM HEPES pH 7.9, 1 M NaCl, 0.5 mM MgCl_2_, 1 mM β-mercaptoethanol) over 40 column volumes and analysed by SDS-PAGE. The purest fractions were pooled, concentrated, and polished with a Superdex 200 Increase 10/300 GL column (Cytiva) in Strep Buffer. Peak fractions were pooled, concentrated, and snap-frozen in liquid nitrogen.

WDR82, PNUTS_WBD_ – WDR82, Symplekin_NTD_ – Ssu72, and Symplekin ^DPR-AAA^ – Ssu72 were purified essentially as described for full-length PNUTS except for the following modifications. Cells were resuspended in Strep Buffer supplemented with 2 mM PMSF, 2.4 mL/L cell culture of Biolock (IBA), and EDTA-free protease inhibitor tablets (Roche). Cells were lysed by addition of Triton X-100 to a final concentration of 0.5 % Triton-X 100. Lysate was clarified by centrifugation and applied to 3 mL Streptactin XT 4 Flow High-Capacity resin (IBA) in a gravity flow column. Resin was washed with 10 CVs of Strep Buffer 3 times followed by elution with 50 mM Biotin (IBA) in Strep Buffer. The C-terminal StrepII-tag was cleaved by addition of home-made TEV protease at a 1:20 target protein to TEV protease ratio and incubation at 4°C overnight. Complexes were then purified by anion exchange and size exclusion chromatographies as described for full-length PNUTS. Peak fractions were pooled, concentrated, and either used immediately for crystallisation trials or flash frozen in liquid nitrogen.

To reconstitute the PNUTS_WBD_ – WDR82 – Symplekin_NTD_ – Ssu72 complex for crystallisation, PNUTS_WBD_ – WDR82 and Symplekin_NTD_ – Ssu72 dimers were expressed and purified separately by Streptactin affinity and anion exchange chromatography as described above. Subcomplexes purified from anion exchange chromatography were concentrated and mixed with PNUTS_WBD_ – WDR82 at a slight molar excess (molar ratio of 1:1.2) for 15 minutes on ice in Strep Buffer followed by purification on a a Superdex 200 Increase 10 /300 GL column equilibrated in the same buffer. Peak fractions showing stoichiometric amounts of all subunits by SDS-PAGE were pooled, concentrated to ∼ 17 mg/mL, and used immediately for crystallisation trials.

To prepare the 10-subunit CPA, cells were co-infected with two baculoviruses, one harbouring genes for the Polymerase Module (CPSF160/Cft1, CPSF30/Yth1, FIP1L1/Iss1, WDR33/Pfs2-STREP tag and PAPOLA/Pla1) and another harbouring genes for the Nuclease and Phosphatase Modules without PNUTS, WDR82, and PP1 (Ysh1, Cft2, Pta1-STREP, Ctf1, Ssu72). Baculoviruses were infected at a 1:1 ratio (v:v) relative to each other and for a total virus:culture ratio of 1:50 (v:v). Cells were harvested after 72 hours post-infection and resuspended in Strep Buffer. CPA was purified by Streptactin affinity and anion exchange chromatographies as described for full-length PNUTS except that home-made TEV protease was added to the Streptactin affinity eluate at a 1:20 ratio (w:w) and the twin-StrepII tags on Symplekin and WDR33 were cleaved overnight at 4°C. During anion exchange chromatography, the polyA polymerase Pla1 (PAPOLA in human) dissociates and elutes as a distinct peak at lower salt. Fractions corresponding to Pla1 and the rest of core CPF were pooled and the salt concentration diluted back down to 150 mM. The complex was then concentrated to < 100 µL and injected onto a Superose 6 Increase 3.2/300 GL column (Cytiva) equilibrated in Strep Buffer. Peak fractions were pooled, concentrated, and flash-frozen in liquid nitrogen.

### Crystallisation and structure determination

#### PNUTS_WBR_ – WDR82

PNUTS_WBR_ – WDR82 at 9.85 mg/mL was crystallised using the sitting-drop vapour diffusion set-up at 20°C. Crystals used for structure solution appeared overnight in two conditions: the first condition was 0.05 M Calcium chloride dihydrate, 0.1 M MES monohydrate pH 6.0, 45% v/v Polyethylene glycol 200, while the second condition was 10% w/v PEG 8000, 20% v/v ethylene glycol 0.03 M of each ethylene glycol 0.1M MES/imidazole pH 6.5. Crystals reached full size after a week and were harvested without addition of cryoprotectant before flash freezing in liquid nitrogen. X-ray diffraction data were collected on the I04 beamline at Diamond Light Source (Harwell, UK). Data collection and refinement statistics are summarised in Supplementary Table 2. Data were automatically processed with autoPROC/STARANISO^125^. The structure was solved via molecular replacement in PHASER^126^, using an AlphaFold2 generated model of WDR82 as the initial search model^127^. PNUTS_WBR_ was built manually into the electron density in COOT^128^.

#### PNUTS_WBR_ – WDR82 – Symplekin_NTD_ – Ssu72

PNUTS_WBR (565 – 644)_ – WDR82 – Symplekin_1 – 307_ – Ssu72 at 17 mg/mL was crystallised using the sitting-drop vapour diffusion set-up at 20°C. Small crystals appeared in a condition containing 100 mM MOPS 7.5, 100 mM Magnesium Acetate, and 12 % PEG 8,000 after 9 days. The drop was composed of 200 nL protein and 100 nL reservoir solution. The crystals were resuspended in 50 μL of the reservoir solution and crushed by vortexing 3 times for 30 seconds using a plastic bead. The seed stock was diluted 1:2 in reservoir solution and used for microseed sparse-matrix screening in sitting-drop vapour diffusion set-up at 20°C. Crystals appeared after 2 – 4 days in multiple conditions and grew to maximal size after 10 – 14 days. Crystals were cryoprotected with 30 % glycerol before flash freezing in liquid nitrogen. X-ray diffraction data were collected on the I04 beamline at Diamond Light Source (Harwell, UK). Data collection and refinement statistics are summarised in Supplementary Table 2. Data were automatically processed with autoPROC/STARANISO. The structure was solved via molecular replacement in PHASER. The input models were the experimentally-solved crystal structure of PNUTS_WBR_ – WDR82 (PDB: 9SCE)) and Alphafold2 generated models of *S*. *pombe* Symplekin_NTD_ and Ssu72. The model was iteratively refined through cycles of refinement in PHENIX followed by manual rebuilding in COOT.

### Analytical size exclusion chromatography (SEC)

Small-scale reconstitution experiments were performed in total reaction volumes of 50 μL in STREP Buffer (50 mM HEPES pH 7.9, 150 mM NaCl, 0.5 mM MgCl_2_, 1 mM β-mercaptoethanol). Individually-purified WDR82, Symplekin_NTD_ – Ssu72, Symplekin ^DPR–AAA^– Ssu72, PNUTS_WBD_ –WDR82 as described above were mixed in the combinations described for final protein concentrations of 64.2 μM in 50 μL total reaction volumes and incubated for 15 minutes on ice. Reactions were then injected onto a Superdex 200 Increase 5/150 GL column equilibrated in STREP Buffer (Cytiva) using a 100 μL loop.

### XL-MS of PNUTS – WDR82 – Symplekin – Ssu72

For crosslinking coupled to mass spectrometry, 10 μg PPW-Symplekin-Ssu72 from size exclusion chromatography at 0.5 mg/mL was crosslinked with 8 mM BS3 on ice for 1 hour. Samples were then quenched with 100 mM Tris pH 7.5. Protein was denatured incubating with 4 M Urea for 10 mins at room temperature, 675 rpm shaking. Samples were then incubated with 10 mM TCEP (Tris (2-carboxyethyl) phosphine) from 30 minutes at room temperature to reduce cysteines. Sample was then incubated with 50mM chloroacetamide (CIAM) for 30 minutes in the dark at room temperature samples were then pre-digested using LysC (1 μg LysC per 100 μg sample) for 2 hrs at 37°C shaking at 800 rpm. 100 mM Ammonium Bicarbonate was then added to the sample to dilute urea to 2 M. Calcium chloride was then added to 2 mM final concentration and the sample was then digested with trypsin at a ratio of 1 μg per 40 μg sample, incubated for 16 – 20 hours at 37 °C shaking at 800 rpm. Trypsin digestion was quenched with 5% formic acid, stored at -20 °C until analysis.

### LC-MS/MS analysis

Dried peptides were resuspended into 5% acetonitrile / 5% formic acid before LC-MS/MS analysis. Peptides were separated by nano liquid chromatography (Ultimate RSLC 3000) coupled in line a QExactive mass spectrometer equipped with an Easy-Spray source (Thermo Fischer Scientific). Peptides were trapped onto a C18 PepMac100 precolumn (300µm i.d.x5mm, 100Å, ThermoFischer Scientific) using Solvent A (0.1% Formic acid, HPLC grade water). The peptides were further separated onto an Easy-Spray RSLC C18 column (75um i.d., 50cm length, Thermo Fischer Scientific) using a 30 (Symplekin^WT^ and Symplekin ^DPR-AAA^ and mutant), 60 (PNUTS-FTP-TANDEM), 120 (PPW-Symplekin-Crosslinking) and 180 mins (PPW-STABLE-COMPLEX) linear gradient (15% to 35% solvent B (0.1% formic acid in acetonitrile)) at a flow rate 200nl/min. The raw data were acquired on the mass spectrometer in a data-dependent acquisition mode (DDA). Full-scan MS spectra were acquired in the Orbitrap (Scan range 350-1500m/z, resolution 70,000; AGC target, 3e6, maximum injection time, 50ms. For 30/60 mins and 120/180mins gradients, the 10 and 20 most intense peaks were selected respectively for higher-energy collision dissociation (HCD) fragmentation at 30% of normalized collision energy. HCD spectra were acquired in the Orbitrap at resolution 17,500, AGC target 5e4, maximum injection time of 120ms with fixed mass at 180m/z. Charge exclusion was selected for unassigned and 1+ ions. The dynamic exclusion was set to 5s, 20s, 40s and 60s for 30-, 60-, 120- and 180-mins gradients respectively.

### Mass spectrometry data processing

For protein identification of FLAG-PNUTS (from Ppn1-FTP) pulldown samples, tandem mass (MS/MS) spectra were searched using Sequest HT in Proteome discoverer software version 1.4 against a protein sequence database containing 5,507 protein entries, including 5,224 *S. pombe* proteins (Uniprot release from 21-11-23) in which the PNUTS protein sequence was replaced by user defined FLAG-tagged PNUTS protein sequence and 283 common contaminants. During database searching cysteines (C) were considered to be fully carbamidomethylated (+57,0215, statically added), methionine (M) to be fully oxidised (+15,9949, dynamically added), all N-terminal residues to be acetylated (+42,0106, dynamically added). Two missed cleavages were permitted. Peptide mass tolerance was set at 50ppm on the precursor and 0.6 Da on the fragment ions. Data was filtered at FDR below 1% at PSM level. For cross-linked peptides identification, tandem mass spectra were searched using pLink software version 2.3.9 against a protein sequence database containing *Spodoptera frugiperda* protein sequences (Uniprot release from 21-02-26) combined with user defined protein sequences namely tagged protein sequences of interest, and 283 common contaminants. During pLink searches, BS3 crosslinker was selected, cysteines (C) were considered to be fully carbamidomethylated (+57,0215, statically added), methionine (M) to be fully oxidised (+15,9949, dynamically added), all N-terminal residues to be acetylated (+42,0106, dynamically added). Three missed cleavages were permitted. Peptide mass tolerance was set at 20ppm on the precursor and fragment ions. Data was filtered at False Discovery Rate (FDR) below 5% at PSM level. The result of the mass spectrometry analyses are summarised in Supplementary Table 3.

### *In vitro* dephosphorylation assay

We purified recombinant PNUTS_420-710_, comprising the PP1- and WDR82-binding regions, WDR82, and PP1 (Figure S1D), and tested the effect of PNUTS_420-710_ and WDR82 on PP1 activity. As substrates, we employed either Spt5 constructs containing KOW5 with seven CTR repeats (KOW_5_-CTR_7_) *in vitro* phosphorylated by recombinantly reconstituted *S. pombe* CDK9/Cyclin T1^32,43^ or short phospho-CTR-peptides (1 and 4 repeats). PP1 alone (4 μM), or PP1 (4 μM) mixed with PNUTS_420-710_ (8 μM) with or without WDR82 (8 μM), was pre-equilibrated at 25 °C for 15 min. Next, samples were diluted in Reaction Buffer (20 mM HEPES pH 7.5, 100 mM NaCl, 2 mM MnCl₂, 1 mM β-mercaptoethanol) to final concentrations of 20 nM PP1 and 40 nM PNUTS_420-710_ or WDR82. Dephosphorylation reactions were performed at 25 °C and initiated by addition of CTR_1_, CTR_4_, or Spt5 Kow_5_–CTR_7_ substrates at final concentrations of 80 μM, 20 μM, and 12 μM, respectively. At designated time points, reactions were quenched with Malachite Green working reagent (Sigma-Aldrich). Phosphate release was quantified by measuring absorbance at 620 nm on a PHERAstar FS plate reader (BMG Labtech) or a DeNovix spectrophotometer and converting values to concentrations using a standard curve generated from known amounts of inorganic phosphate. Negative values were offset to zero. Measurements deviating by more than 100% from the mean of replicate samples at the same time point were classified as outliers and excluded. Curves were fitted using non-linear least squares regression in R (v4.3.1) with the nls function; for reactions using the Kow_5_–CTR_7_ substrate, where nls convergence was not achieved, a LOESS regression (loess function) was applied. Statistical differences among groups at the 60-min time point were assessed using the Kruskal–Wallis rank-sum test, followed by Dunn’s post hoc test with Bonferroni correction when appropriate. All analyses were performed in R environment.

### Pulldown assays

For pulldown assay to test for interaction between core CPA and PPW complex all proteins or protein complexes were recombinantly expressed and purified from Sf9 cells, except for PP1, which was purified from yeast cells (via FLAG tag). Co-expression of 10 CPA subunits in Sf9 cells was performed using two MultiBac constructs.

The assays were performed in STREP Buffer (50 mM HEPES pH 7.9, 150 mM NaCl, 0.5 mM MgCl_2_, 1 mM β-mercaptoethanol). 0.5 μM twin-STREPII tagged PNUTS or PNUTS-WDR82 complex was mixed with 1 μM CPA, PP1-3xFLAG, and/or WDR82 in 50 μL total volume. 20 μL Streptactin XT high-capacity resin (IBA Life Sciences) equilibrated in Strep Buffer was used per pulldown and the mixture was incubated at 4 °C for 1 hour. Resin was washed twice with 200 μL Strep Buffer supplemented with 0.05 % Triton X-100. Bound protein was eluted by boiling the resin in 30 μL Laemmli Buffer. Elutions were split in half and loaded onto separate NuPAGE 4 - 12 % gels (Thermofisher). Each gel was either stained with InstantBlue Coomassie Protein Stain (Abcam) or Western Blotted using HRP-Conjugated Anti-FLAG M2 Antibody (Sigma-Aldrich, # A8592). Western blots were developed using Thermo Scientific™ Pierce™ ECL Western Blotting Substrate and imaged in a Molecular Imager Gel Doc XR System (Biorad).

### TT-seq and Bioinformatic Analyses

Cultures were grown in YES medium (uracil 10 mg/L) to OD_600_ ≈ 0.5. TT-seq was performed in duplicates following published procedures with minor adjustments^129^. Cultures were grown in YES medium (uracil 10 mg/L) either at 25 °C (for degron depletion experiments) or 30°C to OD_600_ ≈ 0.5. Cells were pulse-labelled with 5 mM 4-tU for 6 min, harvested, and snap-frozen. Before RNA extraction, *S. pombe* samples were normalised by OD and supplemented with a 4-tU-labelled *S. cerevisiae* spike-in (100:1). RNA was isolated using the hot-phenol method, treated with DNase, fragmented with NaOH, and conjugated to an MTSEA-biotin-XX linker. Biotinylated RNA was purified with the µMACS streptavidin kit, and fragment sizes were assessed using a Bioanalyzer. Sequencing libraries were generated using the NEBNext Ultra II Directional RNA Library Prep Kit (Illumina) and sequenced on a NextSeq 500. Reads were processed with fastp^130^ and aligned to a combined *S. pombe*/*S. cerevisiae* reference using STAR^131^. Species-specific reads were separated by chromosome identifiers, and uniquely mapped *S. cerevisiae* reads were obtained with SAMtools^132^ for spike-in normalisation. Metagene analyses and heatmaps were generated using deepTools (log_2_ ratios, gene-body scaling when indicated), restricting analyses to ∼3200 coding genes lacking same-strand neighbours within 250 bp of the TSS or PAS. Changes in 3’UTR usage in phosphatase mutants (*pp1*Δ, *pp1*^R245Q^, *wdr82*Δ, *pnuts*Δ, *pnuts*^W510A^) or Symplekin complementation strains were analysed from spike-in normalised BigWig files. 3’ UTR expression was quantified as the ratio of UTR to gene body signal density for ∼4,489 protein-coding genes with 3’ UTRs ≥50 bp. Genes with ≥1.5-fold increase in ≥2 mutants (514 genes) or Symplekin^DPR-AAA^ vs WT complementation strains (526 genes) were classified as upregulated. Hexamer enrichment in upregulated gene 3’ UTRs was assessed by hypergeometric test with Benjamini-Hochberg FDR correction (significant: FDR < 0.05, |log₂FC| > 1). Transcription termination defects were quantified as increased signal in a 100 bp post-TES window relative to gene body density in 3,663 protein-coding genes with 3’ UTRs ≥50 bp, expression ≥10, and without gene on the same strand within 250bp between mutant and WT. Overlap between multi-mutant phosphatases (defects in ≥2 mutants) and Symplekin mutant/WT was evaluated using Venn diagrams and heatmaps showing pairwise percentage overlaps of the smallest set.

## DATA AVAILABILITY

The model and map for the structures presented in this study have been deposited in the PDB database with the identifiers 9SCE (PNUTS_WBR_ bound to WDR82, crystal 1) and 9SYQ (Symplekin_NTD_ - Ssu72 - PNUTS_WBR_ – WDR82 complex - crystal form 1). TT-seq data have been deposited in NCBI’s Gene Expression Omnibus and are accessible through GEO Series accession number GSE308656.

## ACKNOWLEDGEMENTS

We thank members of the Vasilieva lab for helpful discussions, advice, and valuable comments on the manuscript. This work was supported by the Wellcome Trust Senior Fellowship and BBSRC grants to L.V. (WT106994/Z/15/Z, BB/Y00194X/1 and BB/Y004590/1), the Wellcome Trust Investigator Award to J.M.G. (200835/Z/16/Z and 222510/Z/21/Z), Wellcome Trust studentship to A.A., and the Emmy Noether Programme of the Deutsche Forschungsgemeinschaft (DFG) to C.K. (KI 1657/2-1). S.S.H. is funded by a scholarship grant (MM1/23/ID:1563097) from the Egyptian Mission Department.

## AUTHOR CONTRIBUTIONS STATEMENT

A.A., K.K., A.S. and L.V. conceived and designed the experiments. A.A. cloned baculovirus constructs, generated the original bacterial PP1 expression construct, and purified native PP1 and recombinant proteins from insect cells. A.A., E.B.A. and J.M.G. solved the crystal structures of PNUTS-WDR82 and PNUTS-WDR82-Symplekin-Ssu72 complexes. A.A. and C.H. purified the native cleavage and polyadenylation complex and assessed Pol II and Spt5 phosphorylation status in phosphatase mutant cells. K.K. designed and generated some baculovirus constructs, performed TT-seq and data analysis for phosphatase mutants, and contributed to generating the Symplekin mutant strain. A.S. generated Symplekin mutant strains, initial bacterial PNUTS expression constructs, and performed TT-seq experiments and analyses in these strains. N.S. optimised bacterial expression and purification of recombinant PP1 and PNUTS and established and performed phosphatase assays. E.A. and C.K. performed microscopy experiments and quantification of PNUTS cellular localisation. M.F. performed mass spectrometry experiments and analysis. S.S.H. performed Western blot analyses of Pol II phosphorylation in Symplekin strains. A.A., K.K., A.S. and L.V. prepared the figures and wrote the manuscript. All authors read and commented on the manuscript.

## COMPETING INTERESTS STATEMENT

The authors declare no competing interests.

## Supplementary Figures

**Figure S1**

A) Nuclear localisation of *S. pombe* PNUTS. Immunofluorescence against FLAG in untagged WT and PNUTS-FTP cells. The merged channel shows FLAG in green and DAPI in magenta. Scale bar = 5 µm. Bottom: Quantitation of FLAG signal (n = 2 independent experiments). Black horizontal lines mark the median and the thick grey line the mean FLAG signal ratio of the untagged strain.

B) Purification of PPW subunits for the phosphatase assay. PNUTS_420–710_ was purified from *E. coli* by sequential affinity steps and SEC (lane 1). PP1 was purified from *E. coli* using an intein-based system (lane 2). WDR82-STREP was purified from Sf9 insect cells using Strep-Trap HP chromatography (lane 3).

C) Phos-tag gels of Kow_5_-CTR_7_ phosphorylated by incubation with Cdk9-Pch1.

D) Schematic showing phosphorylated peptide substrate sequences.

E) Phosphatase assay using Kow_5_-CTR_7_ as a substrate.

F) Phosphatase assay using 1× phosphorylated peptide substrate (Spt5-CTR_1_)

**Figure S2**

A) Transcriptional readthrough upon loss of PP1, its catalytic activity or regulatory subunits (WDR82, PNUTS or complementation strain with PNUTS^W510A^ which losses interaction with PP1). Uncropped TT-seq signal in indicated mutants - related to Figure 1B).

B) Changes in different classes of non-coding RNAs in mutants described in (A).

C) Distribution of 3’UTR lengths which were upregulated at least 1.5-fold in at least 2 mutants described in (A), n=514 compared to 3’UTRs that has not been assigned to be changed in any of the mutants (n=3419).

D) Comparison of 3’UTRs that were upregulated (as in C) to control.

E) Hexamer enrichment analysis for sequences found in upregulated 3’UTRs versus the unchanged set (as in C-D).

**Figure S3**

A) PNUTS and WDR82 were co-expressed in insect cells and analysed by size-exclusion chromatography, revealing co-elution as a stable heterodimeric complex.

B) Purified PNUTS_WBR_-WDR82 and Symplekin_NTD_-Ssu72 heterodimers were combined and resolved by size-exclusion chromatography, yielding a stable reconstituted hetero-tetrameric complex.

C) Electron density encapsulating Ssu72 and Symplekin_NTD_ in tetramer structure. Only part of Ssu72 is present in crystal structure corresponding to LMWPTP domain. Partial Ssu72 model is overlayed with AlphaFold predicted structure.

D) A Ssu72 protein topology diagram highlighting CAP and LMWPTP domains. Tetrameric structure only contained density covering part of LMWPTP domain.

E) Ssu72 interaction with Symplekin is conserved in yeast and human. Structural superimposition of PNUTS_WBR_–WDR82-Symplekin_NTD_-Ssu72 (this work) with Syplekin-Ssu72 (human)^78^.

F) The PNUTS_WBR_–WDR82 heterodimer maintains the same overall conformation in PNUTS_WBR_–WDR82-Symplekin_NTD_-Ssu72 tetramer as observed when PNUTS_WBR_–WDR82 structure solved independently.

**Figure S4**

A) Electron density map covering the interface between the Symplekin DPR motif and the WDR82 binding pocket.

B) Analyses of the complex for WDR82 with Symplekin_NTD_ –Ssu72 or Symplekin_NTD_^DPR-AAA^-Ssu72 formation by analytical SEC (Superdex200). Symplekin_NTD_^DPR-AAA^ mutant formed a heterodimer with Ssu72, however, failed to assemble WDR82 into the hetero-trimer, in contrast to wild-type SymplekinNTD –Ssu72.

C) Domain maps *S. pombe* PNUTS and PNUTS_WPL-G/A_.

D) PNUTS_WPL-G/A_ mutant retained PP1 binding but failed to pull down Symplekin_NTD_–Ssu72, even with WDR82 present, highlighting the importance of the WDR82-interacting surface in complex assembly.

**Figure S5**

A) Growth assay indicating that depletion of Symplekin-AID result in lethality when grown on auxin plates.

B) Depletion of Symplekin-AID following treatment with 1 mM Auxin. Ponceau stain used as loading control.

C) Overlap between read-through observed in the PPW mutants with the effect observed in Symplekin^DPR-AAA^

D) Distribution of 3’UTR lengths which were upregulated at least 1.5-fold in Symplekin^DPR-AAA^ compared to Symplekin^WT^, n=526 compared to 3’UTRs that has not been assigned to be changed in control (n=3935).

E) Comparison of CG content in 3’UTRs that were upregulated in Symplekin^DPR-AAA^ compared to Symplekin^WT^.

F) Changes in different classes of non-coding RNAs in Symplekin^WT^ and Symplekin^DPR-AAA^

**Figure S6**

A) Multiple sequence alignment reveals strong conservation within the WDR82 region that forms the pocket engaging DPR motifs, including those found in Symplekin.

B) Comparative domain organization of Symplekin from multiple organisms, combined with sequence alignment, reveals that the DPR motif is uniquely present in *S. pombe*.

C) AlphaFold predictions of Symplekin NTD structures from various organisms demonstrate that the loop between helices 7 and 8 varies substantially across species and is not evolutionarily conserved.

D) Superimposing the *S. pombe* Symplekin NTD structure with the predicted *S. cerevisiae* NTD reveals substantial divergence, implying that, in budding yeast, the connection between CPA and the phosphatase module is likely mediated by an alternative protein (like Pti1).

E) AlphaFold predictions of the interactions within ZC3H4-WDR82-PNUTS complex. DPR is provided by ZC3H4 whereas WPL/L and IL-like by PNUTS.

F) Predicted ZC3H4-DPR-WDR82 binding interfecate.

G) Structural prediction of WDR82 in complex with SET1 in *H. sapiens*.

## Notes

### Competing Interest Statement

The authors have declared no competing interest.

