## Supplementary figures for "The PP1 phosphatase complex coordinates pre-mRNA 3’end processing and termination of RNA polymerase II transcription"

Figure S1

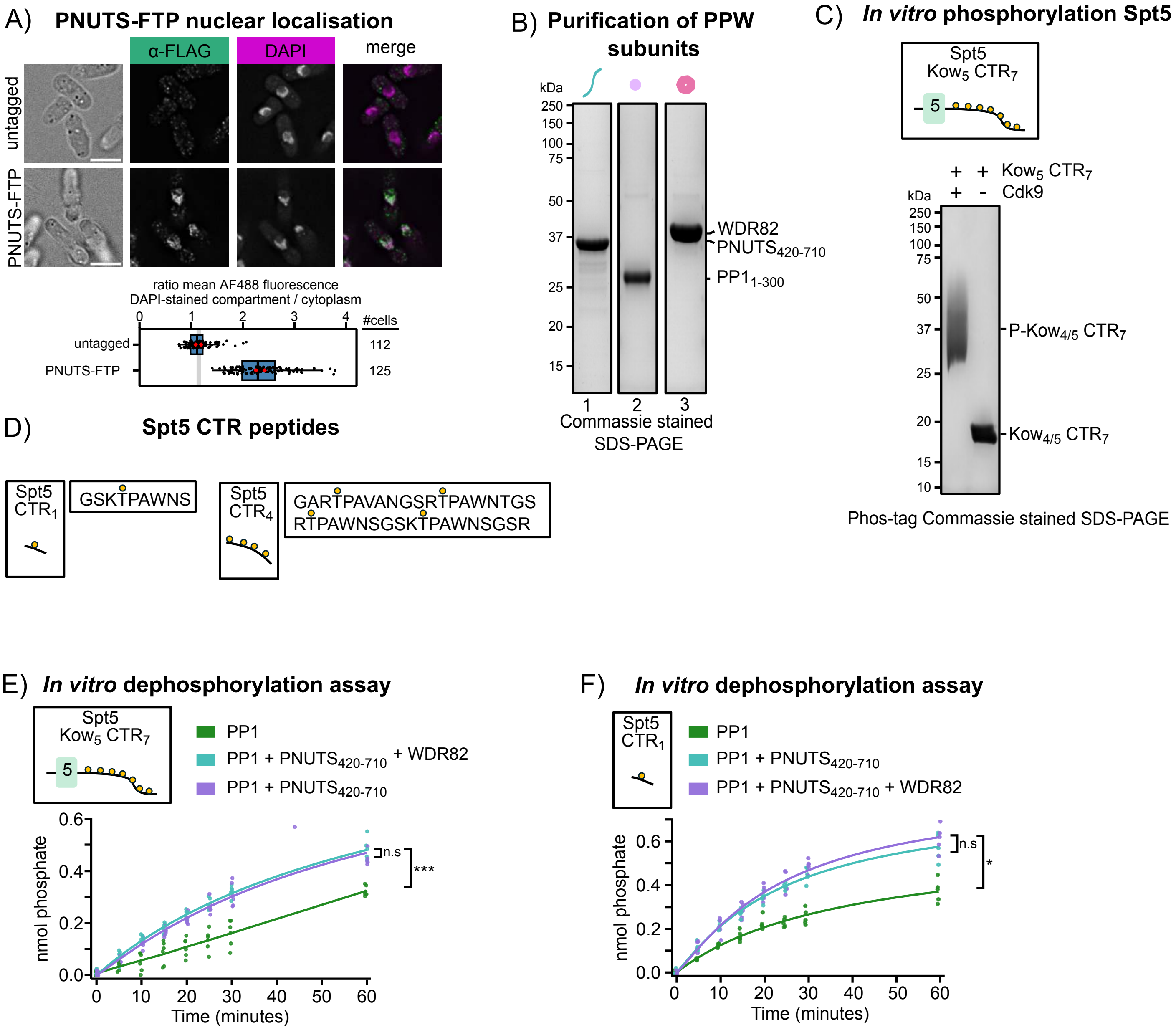
A)
PNUTS-FTP nuclear localisation
α-FLAG
DAPI
merge
untagged
PNUTS-FTP
ratio mean AF488 fluorescence
DAPI-stained compartment / cytoplasm
0
1
2
3
4
#cells
untagged
PNUTS-FTP
112
125
B)
Purification of PPW subunits
kDa
250
150
100
75
50
37
25
20
15
1
2
3
WDR82
PNUTS420-710
PP11-300
Commassie stained
SDS-PAGE
C)
In vitro phosphorylation Spt5
Spt5
Kow5 CTR7
5
+ + Kow5 CTR7
+ - Cdk9
kDa
250
150
100
75
50
37
25
20
15
10
P-Kow4/5 CTR7
Kow4/5 CTR7
Phos-tag Commassie stained SDS-PAGE
D)
Spt5 CTR peptides
Spt5
CTR1
GSKTPAWNS
Spt5
CTR4
GARTPAVANGSRTPAWNTGS
RTPAWNSGSKTPAWNSGSR
E)
In vitro dephosphorylation assay
Spt5
Kow5 CTR7
5
PP1
PP1 + PNUTS420-710 + WDR82
PP1 + PNUTS420-710
nmol phosphate
0.0
0.2
0.4
0.6
n.s
***
0
10
20
30
40
50
60
Time (minutes)
F)
In vitro dephosphorylation assay
Spt5
CTR1
PP1
PP1 + PNUTS420-710
PP1 + PNUTS420-710 + WDR82
nmol phosphate
0.0
0.2
0.4
0.6
n.s
*
0
10
20
30
40
50
60
Time (minutes)

### Figure S2

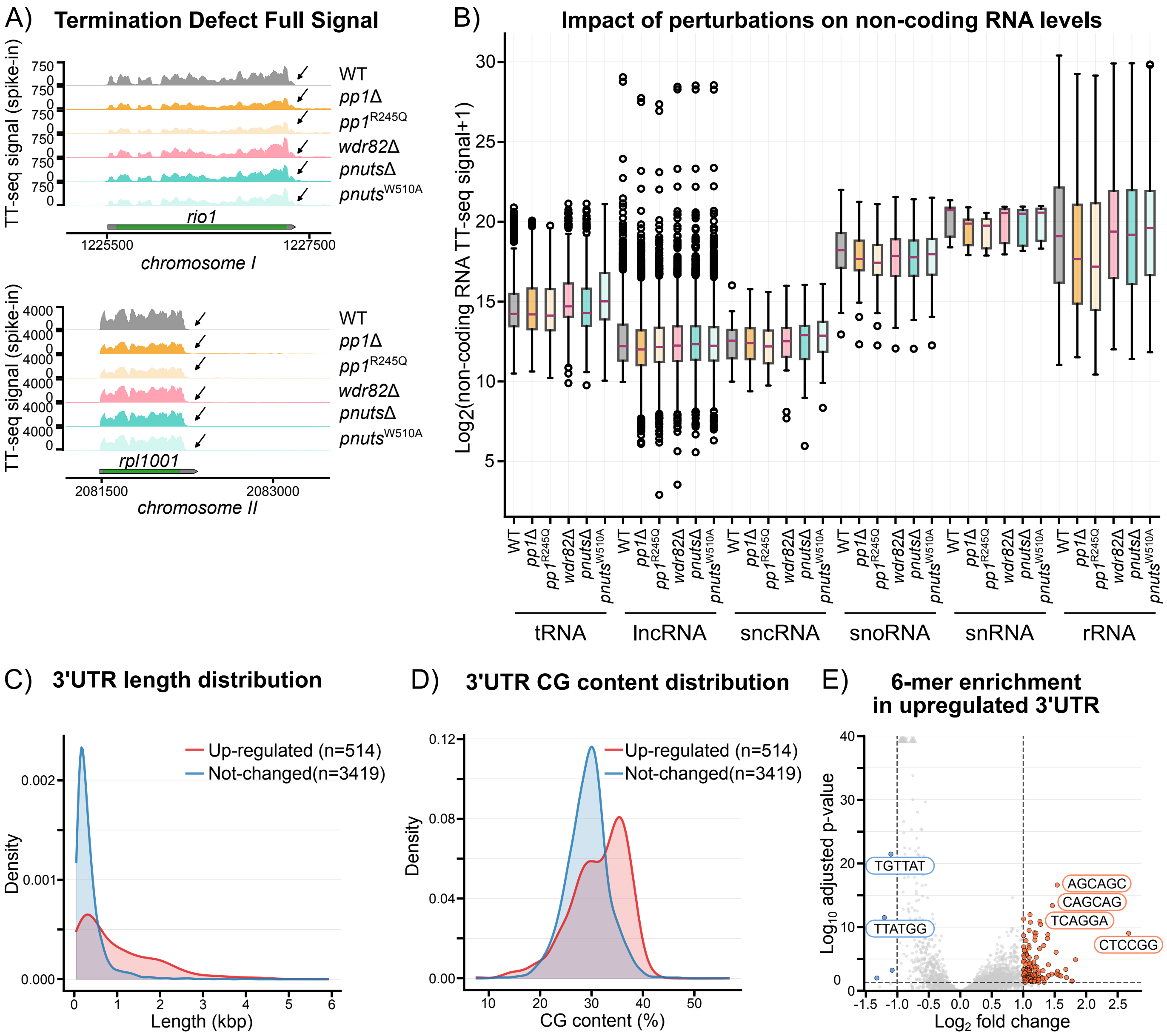

A) Termination Defect Full Signal
TT-seq signal (spike-in)
WT
pp1Δ
pp1R245Q
wdr82Δ
pnutsΔ
pnutsW510A
rio1
1225500
1227500
chromosome I
rpl1001
2081500
2083000
chromosome II
B) Impact of perturbations on non-coding RNA levels
Log2(non-coding RNA TT-seq signal+1)
tRNA
lncRNA
sncRNA
snoRNA
snRNA
rRNA
C) 3'UTR length distribution
Up-regulated (n=514)
Not-changed(n=3419)
Density
Length (kbp)
D) 3'UTR CG content distribution
Up-regulated (n=514)
Not-changed(n=3419)
Density
CG content (%)
E) 6-mer enrichment in upregulated 3'UTR
Log10 adjusted p-value
Log2 fold change
TGTTAT
TTATGG
AGCAGC
CAGCAG
TCAGGA
CTCCGG

Figure S3

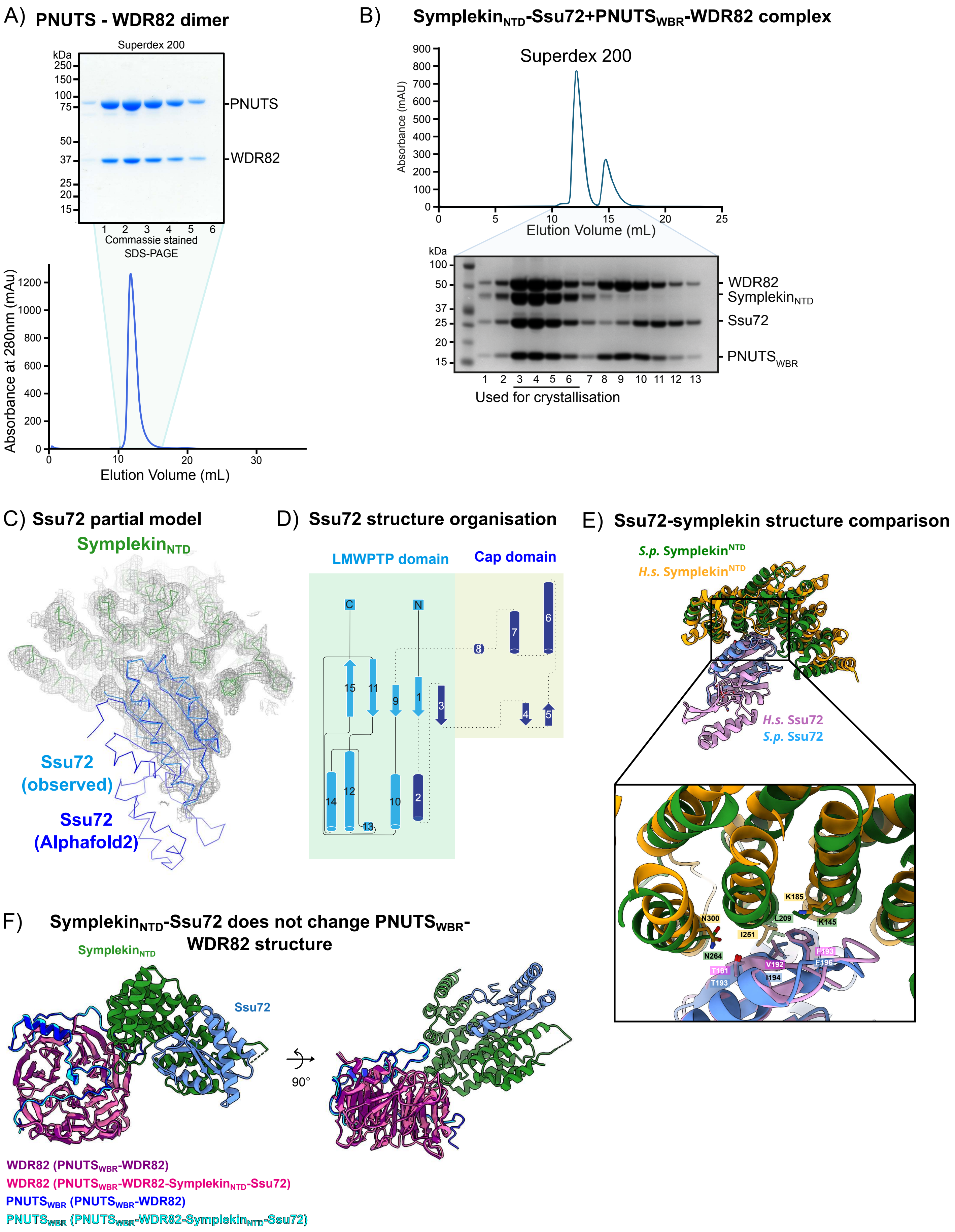
A) PNUTS - WDR82 dimer
Superdex 200
kDa
250
150
100
75
50
37
25
20
15
PNUTS
WDR82
1 2 3 4 5 6
Commassie stained
SDS-PAGE
Absorbance at 280nm (mAu)
1200
1000
800
600
400
200
0
0 10 20 30
Elution Volume (mL)
B) Symplekin$_{NTD}$-Ssu72+PNUTS$_{WBR}$-WDR82 complex
Superdex 200
Absorbance (mAU)
900
800
700
600
500
400
300
200
100
0
0 5 10 15 20 25
Elution Volume (mL)
kDa
100
50
37
25
20
15
WDR82
Symplekin$_{NTD}$
Ssu72
PNUTS$_{WBR}$
1 2 3 4 5 6 7 8 9 10 11 12 13
Used for crystallisation
C) Ssu72 partial model
Symplekin$_{NTD}$
Ssu72 (observed)
Ssu72 (Alphafold2)
D) Ssu72 structure organisation
LMWPTP domain
Cap domain
C
N
E) Ssu72-symplekin structure comparison
S.p. Symplekin$^{NTD}$
H.s. Symplekin$^{NTD}$
H.s. Ssu72
S.p. Ssu72
K185
N300
L209
K145
I251
N264
F193
V192
F196
T191
I194
T193
F) Symplekin$_{NTD}$-Ssu72 does not change PNUTS$_{WBR}$-WDR82 structure
Symplekin$_{NTD}$
Ssu72
90°
WDR82 (PNUTS$_{WBR}$-WDR82)
WDR82 (PNUTS$_{WBR}$-WDR82-Symplekin$_{NTD}$-Ssu72)
PNUTS$_{WBR}$ (PNUTS$_{WBR}$-WDR82)
PNUTS$_{WBR}$ (PNUTS$_{WBR}$-WDR82-Symplekin$_{NTD}$-Ssu72)

Figure S4

A) **Density around DPR motif**

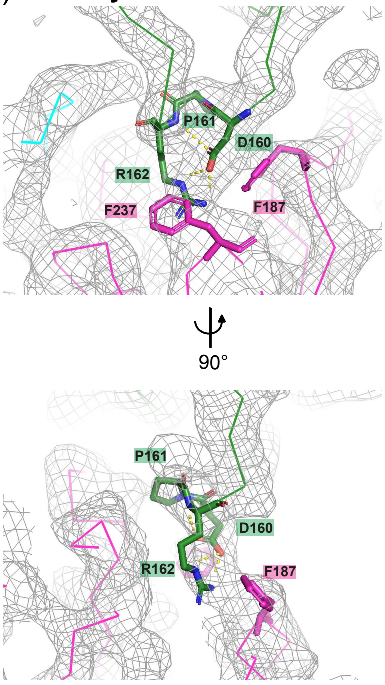


B) **Size Exclusion chromatography**

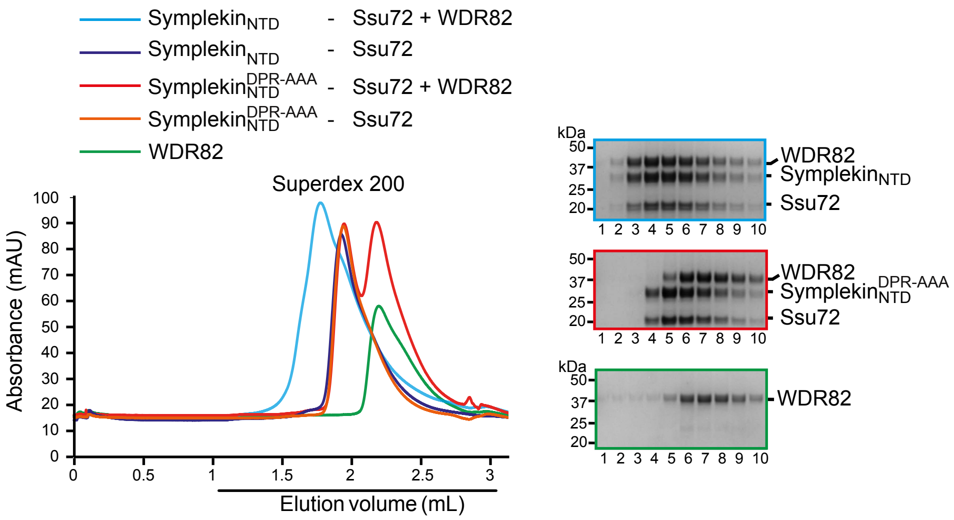


C) **PNUTS constructs used in pulldown assay**

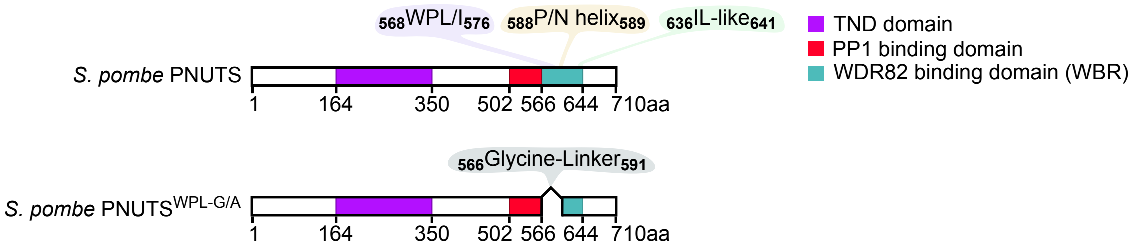


D) **Probing PPW interaction with Symplekin-Ssu72**

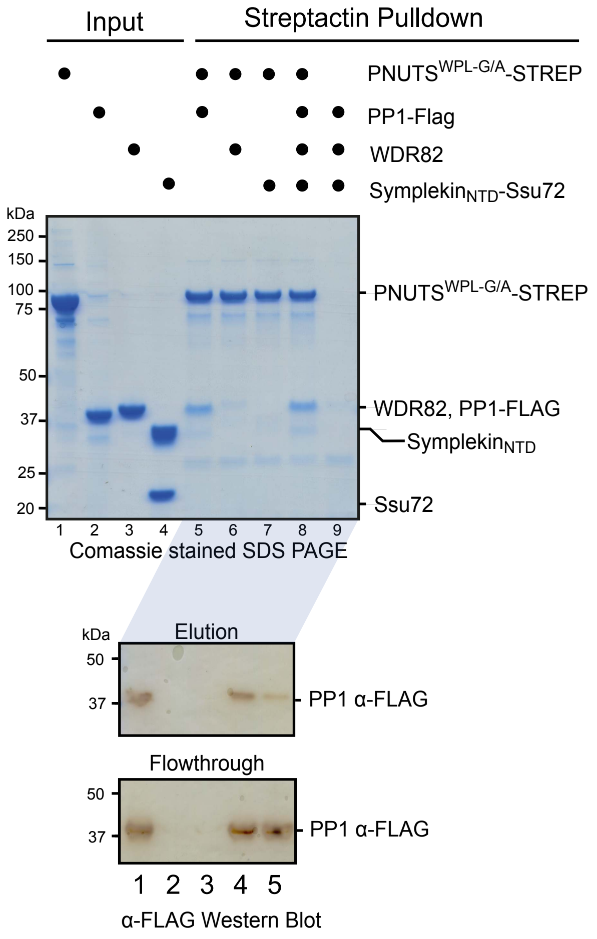

Figure S5

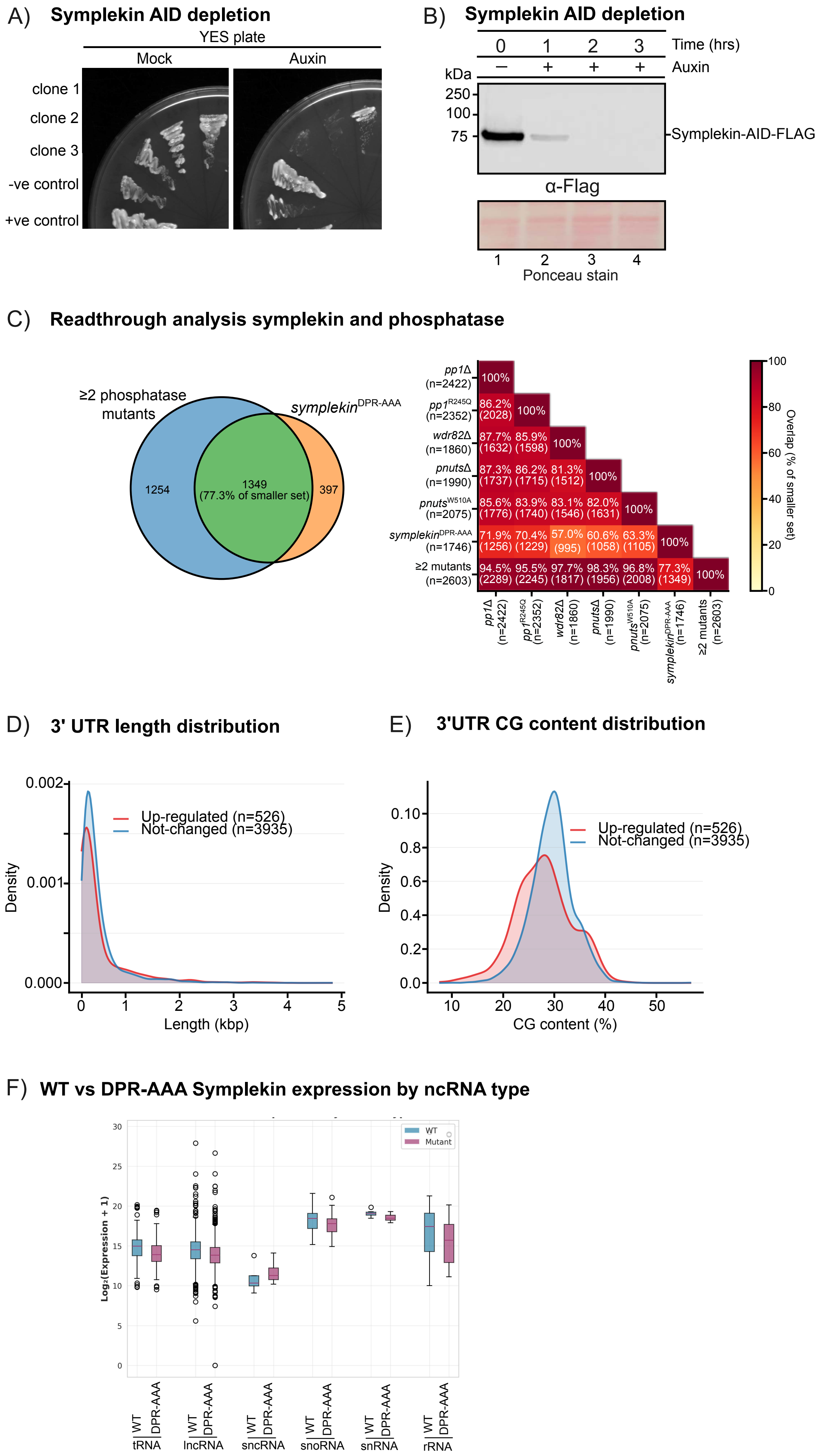
A) Symplekin AID depletion
YES plate
Mock
Auxin
clone 1
clone 2
clone 3
-ve control
+ve control
B) Symplekin AID depletion
0 1 2 3 Time (hrs)
– + + + Auxin
kDa
250
100
75
Symplekin-AID-FLAG
α-Flag
1 2 3 4
Ponceau stain
C) Readthrough analysis symplekin and phosphatase
≥2 phosphatase mutants
symplekin$^{DPR-AAA}$
1254
1349 (77.3% of smaller set)
397
pp1Δ (n=2422)
pp1$^{R245Q}$ (n=2352)
wdr82Δ (n=1860)
pnutsΔ (n=1990)
pnuts$^{W510A}$ (n=2075)
symplekin$^{DPR-AAA}$ (n=1746)
≥2 mutants (n=2603)
100%
86.2% (2028) 100%
87.7% (1632) 85.9% (1598) 100%
87.3% (1737) 86.2% (1715) 81.3% (1512) 100%
85.6% (1776) 83.9% (1740) 83.1% (1546) 82.0% (1631) 100%
71.9% (1256) 70.4% (1229) 57.0% (995) 60.6% (1058) 63.3% (1105) 100%
94.5% (2289) 95.5% (2245) 97.7% (1817) 98.3% (1956) 96.8% (2008) 77.3% (1349) 100%
Overlap (% of smaller set)
0 20 40 60 80 100
D) 3' UTR length distribution
Up-regulated (n=526)
Not-changed (n=3935)
Density
0.000 0.001 0.002
Length (kbp)
0 1 2 3 4 5
E) 3'UTR CG content distribution
Up-regulated (n=526)
Not-changed (n=3935)
Density
0.0 0.2 0.4 0.6 0.8 0.10
CG content (%)
10 20 30 40 50
F) WT vs DPR-AAA Symplekin expression by ncRNA type
WT
Mutant
Log₂(Expression + 1)
0 5 10 15 20 25 30
WT DPR-AAA tRNA
WT DPR-AAA lncRNA
WT DPR-AAA sncRNA
WT DPR-AAA snoRNA
WT DPR-AAA snRNA
WT DPR-AAA rRNA

### Figure S6

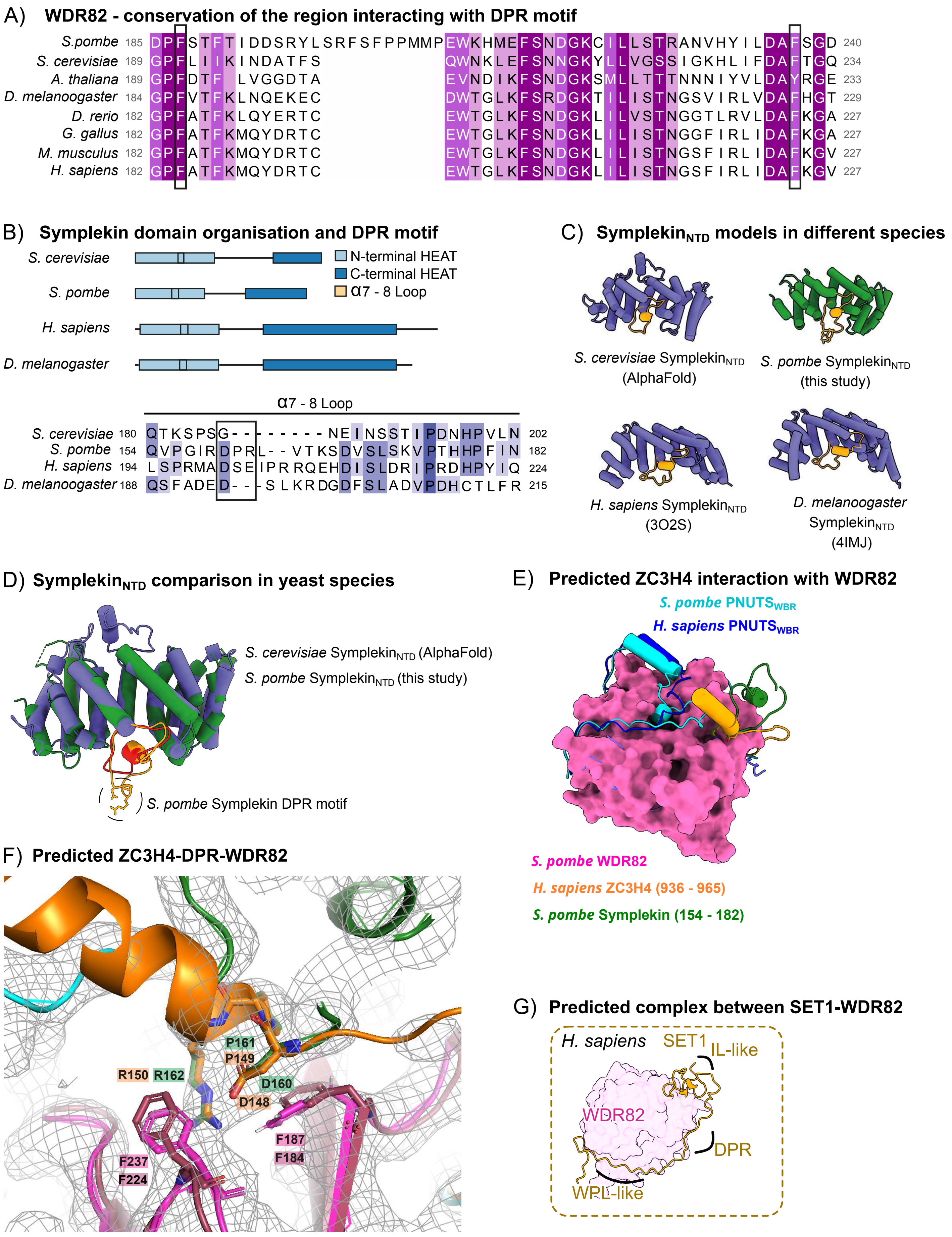
A) WDR82 - conservation of the region interacting with DPR motif
S.pombe 185 DPFSTFTIDDSRYLSRFSFPPMMPEWKHMEFSNDGKCILLSTRANVHYILDAFSGD 240
S. cerevisiae 189 GPFLIIKINDATFS QWNKLEFSNNGKYLLVGSSIGKHLIFDAFTGQ 234
A. thaliana 189 GPFDTF LVGGDTA EVNDIKFSNDGKSMLLTTTNNNIYVLDAYRGE 233
D. melanoogaster 184 GPFVTFKLNQEKEC DWTGLKFSRDGKTILISTNGSVIRLVDAFHGT 229
D. rerio 182 GPFATFKLQYERTC EWTGLKFSNDGKLILVSTNGGTLRVLDAFKGA 227
G. gallus 182 GPFATFKMQYDRTC EWTGLKFSNDGKLILISTNGGFIRLIDAFKGA 227
M. musculus 182 GPFATFKMQYDRTC EWTGLKFSNDGKLILISTNGSFIRLIDAFKGV 227
H. sapiens 182 GPFATFKMQYDRTC EWTGLKFSNDGKLILISTNGSFIRLIDAFKGV 227
B) Symplekin domain organisation and DPR motif
S. cerevisiae
S. pombe
H. sapiens
D. melanogaster
N-terminal HEAT
C-terminal HEAT
α7 - 8 Loop
α7 - 8 Loop
S. cerevisiae 180 QTKSPSG------NEINSSTIPDNHPVLN 202
S. pombe 154 QVPGIRDPRL--VTKSDVSLSKVPTHHPFIN 182
H. sapiens 194 LSPRMADSEIPRRQEHDISLDRIPRDHPYIQ 224
D. melanoogaster 188 QSFADED---SLKRDGDFSLADVPDHCTLFR 215
C) Symplekin$_{NTD}$ models in different species
S. cerevisiae Symplekin$_{NTD}$ (AlphaFold)
S. pombe Symplekin$_{NTD}$ (this study)
H. sapiens Symplekin$_{NTD}$ (3O2S)
D. melanogaster Symplekin$_{NTD}$ (4IMJ)
D) Symplekin$_{NTD}$ comparison in yeast species
S. cerevisiae Symplekin$_{NTD}$ (AlphaFold)
S. pombe Symplekin$_{NTD}$ (this study)
S. pombe Symplekin DPR motif
E) Predicted ZC3H4 interaction with WDR82
S. pombe PNUTS$_{WBR}$
H. sapiens PNUTS$_{WBR}$
S. pombe WDR82
H. sapiens ZC3H4 (936 - 965)
S. pombe Symplekin (154 - 182)
F) Predicted ZC3H4-DPR-WDR82
P161
P149
R150
R162
D160
D148
F187
F184
F237
F224
G) Predicted complex between SET1-WDR82
H. sapiens
SET1
IL-like
WDR82
DPR
WPL-like
